# Continuous contributions of the dorsolateral striatum to movement initiation and execution

**DOI:** 10.1101/2025.11.10.687756

**Authors:** Jiaming Cao, Yujie Zhao, Jianing Yu

**Affiliations:** State Key Laboratory of Membrane Biology, Peking University, 100871, Beĳing, China; School of Life Sciences, Peking University, 100871, Beĳing, China; Academy for Advanced Interdisciplinary Studies, Peking University, 100871, Beĳing, China; IDG/McGovern Institute for Brain Research at Peking University, 100871, Beĳing, China

## Abstract

Spikes in the dorsolateral striatum (DLS) of the basal ganglia correlate with forelimb and whole-body movements, but the extent to which they contribute to movement remains unclear. Using behavior-timed optogenetic inhibition in rats performing a lever-release task, we found that DLS inhibition delayed the initiation of forelimb reaching and conditioned lever release and, when delivered during reaching, interrupted execution. Inhibition during reward retrieval impaired locomotion, especially turning, reducing movement speed and resulting in less efficient trajectories. By contrast, inhibition before the nose poke that triggered reward delivery had little effect, despite substantial DLS activity at the nose poke. These effects were accompanied by bidirectional changes in firing rates in the substantia nigra pars reticulata, a basal ganglia output nucleus, and were much weaker when the dorsomedial striatum was perturbed. Together, these findings suggest that expression of many, but not all, learned movement patterns depends continuously on DLS activity.

**Teaser:** Briefly silencing striatal neurons during behavior reveals that the dorsolateral striatum helps initiate and control the execution of forelimb and whole-body movements in rats.

## Introduction

Behavior has been defined as the total movement produced by an animal (*1*), within which learned movement sequences often involve repeated execution of actions with high precision, low variability, and rapid speed. Despite fine-scale dissections of the brain areas that may contribute to movement generation—including neural circuits in the cortex and basal ganglia (BG)(*2–4*), the roles of different neural structures in producing movements, and whether these roles generalize across movement types, remain unresolved (*5*). The dorsal striatum, the input to the BG, is widely implicated in the learning and execution of procedural skills (*6–8*). Striatal neural activity correlates strongly with behavioral events during procedural tasks and changes with learning (*9–11*). Proposed roles of the striatum include action selection (*12–14*), encoding kinematic parameters of movements (*15*, *16*), controlling movement vigor (i.e., speed and force) (*17–20*), and storing or chunking motor sequences when behaviors become automatic (*9*, *10*, *21*).

Although spikes in the striatum have been functionally correlated with various behaviors, the extent to which these spikes causally influence movement execution remains elusive. In particular, it is not clear whether these spikes contribute to movement on a moment-to-moment basis and whether there is any movement phase (e.g., before initiation or during execution) or type during which striatal spikes become dispensable. Furthermore, prior work using lesions or perturbations has typically focused either on precisely controlled forelimb movements (*8*, *15*, *22–27*) or on whole-body locomotion (*18*, *20*, *27*, *28*). Whether striatal activity contributes uniformly across movement types therefore remains unclear. Moreover, movements that appear continuous at the behavioral level—e.g., moving towards a choice port—can often be decomposed into multiple submovements (*5*, *29–31*), raising the possibility that striatal spikes may contribute continuously to the execution of such actions.

To fill this gap, we performed striatal optogenetic inhibition across multiple phases of a procedural task that involved both forelimb and whole-body movements. We used a simple reaction time (SRT) task in which rats performed a conditioned, tone-cued lever release to obtain a reward (*19*, *32*, *33*). In this task, rats pressed and held a lever, then released it upon hearing a trigger tone. The reward (water) port on the opposite wall of the behavioral chamber required the rat to make a whole-body locomotor movement, including turning, followed by an operant nose poke to obtain a water reward. Both forelimb and whole-body movements were learned operant responses that naturally chain into a motor sequence. First, we characterized how these movement types are represented in the dorsolateral striatum (DLS) using chronic multisite recordings (including Neuropixels probes). Second, by optogenetically activating inhibitory interneurons to suppress striatal projection neurons, we examined how behavior-coupled DLS inhibition affected movement initiation and execution. Finally, by recording from a BG output nucleus, the substantia nigra pars reticulata (SNr), during DLS inhibition, we examined SNr firing properties in the same task and linked the behavioral effects of DLS inhibition to changes in SNr neural activity. Our results demonstrate that event-coupled DLS spikes causally influence the execution of both forelimb and whole-body movements—delaying or interrupting forelimb actions and leading to slower, less efficient whole-body locomotion—whereas comparable perturbations in the nearby dorsomedial striatum (DMS) have only minor effects.

## Results

### Event-locked DLS activity across SRT task phases

Rats were trained to perform an SRT task (*19*, *32–36*). In a single session, the rat moved between a spring-loaded lever and a reward port in a custom behavioral box (Fig. 1a). To start a trial (Fig. 1b), the rat was required to voluntarily initiate a lever press, without any external cue, and hold it for an unpredictable duration (750 or 1500 ms), known as the foreperiod (FP). At the end of the FP, a brief, 0.25-s tone sounded, and the rat released the lever within a 0.6-s response window to obtain a reward. Premature or late responses were not rewarded. The reward port was located on the opposite side of the behavioral box, where a nose poke triggered water drops after a correct lever release. Well-trained rats exhibited stable, reactive lever releases across sessions (Fig. 1c), with low premature responses and a rapid increase in response probability immediately after the stimulus (fig. S1a). Rats formed stereotypical locomotor patterns in the operant box, moving between the reward port and the lever (Fig. 1d). Forelimb reaches were also stereotyped. After training, rats (*n*= 26) released the lever in response to the trigger tone (fig. S1a). The median correct response ratio was 68.8% (interquartile range [IQR]: 65%–76.8%) for short FP trials (750 ms), and 63% (IQR: 55.3%–67.4%) for long FP trials (1500 ms) (fig. S1b). The median reaction time was 0.34 s (IQR: 0.27–0.37 s) for short FP trials and 0.29 s (0.25–0.33 s) for long FP trials (fig. S1c).

**Fig. 1.**
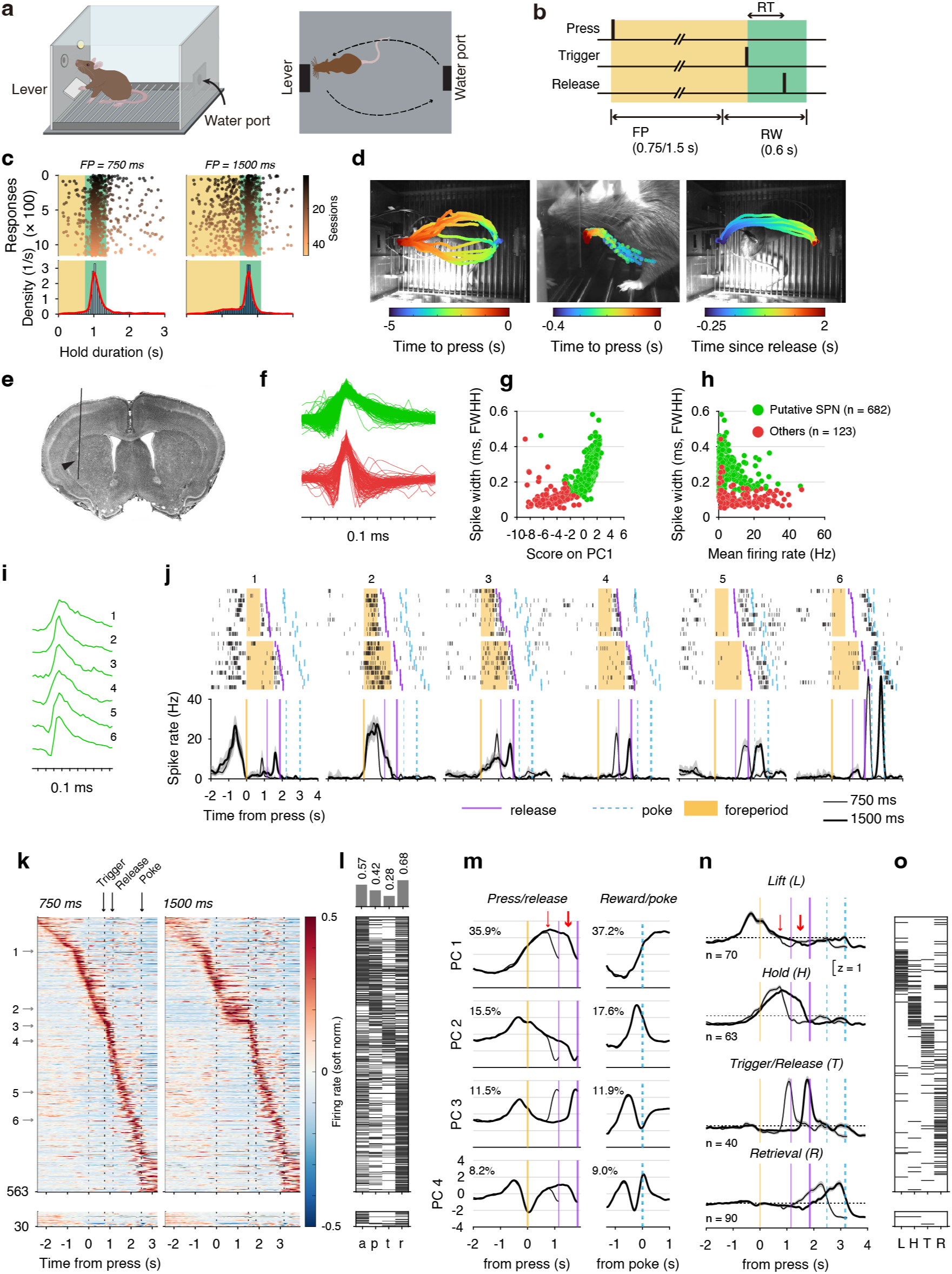
Simple reaction time (SRT) behavior and underlying dorsolateral striatum (DLS) neural dynamics. **a**, Behavioral apparatus schematic. **b**, Task structure (RT: reaction time; FP: foreperiod; RW: response window). **c**, Lever-release responses of a single rat across 47 sessions (25% of trials shown, colored by session index). Top: responses by occurrence time. Bottom: kernel density estimate (red). **d**, Locomotor and forelimb movement trajectories from a single rat in one session. Left: lever approach (top view; frame size: 28×36 cm). Middle: paw reaching for lever presses (side view; frame size: 7×8 cm). Right: reward retrieval (top view). **e**, Example Neuropixels probe track in DLS. **f**, Two spike waveform types in DLS (amplitudes z-scored). **g**, Spike width (full width at half maximum, FWHM) plotted against projection onto the first principal component. Units were classified via unsupervised clustering; green shows putative spiny projection neurons. **h**, Spike width plotted against average firing rate. **i**, Six example waveforms of putative spiny projection neurons. **j**, Spike rasters and spike density functions (SDFs) of examples in **i**. Rows 1–2: Rasters for 10 trials each of 750- and 1500-ms FP (yellow: FP; purple: lever release; cyan: nose poke). Row 3: SDFs by FP type (thin/thick: 750/1500 ms; shading: 95% CI; purple/cyan: median release/poke times). All from one rat’s Neuropixels data. **k**, SDFs of DLS units (rows: units; color: normalized rate). Top/bottom: positively/negatively modulated. Left/right: 750/1500 ms FP. Arrows mark examples from **i–j**. **l**, Modulation by approach (a), press (p), trigger (t) or retrieval (r) for each unit from a GLM (FDR= 0.05). Unit order is the same as in **k**. Histogram showing the proportion of units modulated by each event. **m**, Population projections onto the first four PCs around lever press (left) or port nose poke (right). Thin/thick: 750/1500 ms FP. Colors match those in **j**. Numbers indicate variance explained. Red arrows: trigger onset. **n**, Average SDFs (z-scored) of functional units selected based on their projection loadings (see Materials and Methods). **o**, Index of selected functional units (Lift, Hold, Trigger/Release, or Retrieval); horizontal lines mark unit assignments, ordered from top to bottom to match the colormap order in **k**. Shading denotes SEM in **m** and **n**.

We performed chronic multichannel recordings in the DLS of 10 rats using microwire arrays or silicon probes, of which two were implanted with Neuropixels probes (Fig. 1e). Both wide (0.26 ± 0.07 ms, mean ± SD, *n*= 682) and narrow (0.11 ± 0.05 ms, *n*= 123) waveforms were observed (Fig. 1f–h). We focused our analysis on the wide-waveform units, which likely represent putative spiny projection neurons (SPNs), while the remaining units likely originated from interneurons or *en passant* axons. Putative SPNs in the DLS exhibited low average firing rates (median: 1.27 Hz; IQR: 0.68–2.46 Hz). After excluding units with extremely low firing rates, even when modulated by behavior (maximal event-aligned spike density function [SDF] < 4 Hz), non-significant task-related modulation (as determined by a generalized linear model [GLM], fig. S2a; Equation 1, see Materials and Methods), or substantial inter-spike interval (ISI) violations (> 2% of spikes had ISI < 3 ms), we included 593 units for functional analysis. Most of these units (*n*= 563) exhibited elevated firing during one or more task events (Fig. 1k, l).

During the SRT task, SPN activity was often modulated by one or more behavioral components (Fig. 1i–l), including lever approach/press (example unit 1), lever holding (example units 2, 3), trigger/release (example units 1, 3, 4), and reward retrieval (example units 5, 6).

To capture the modulation of population activity by behavior, we assembled a pseudopopulation using the soft-normalized average firing rate of 593 units and performed principal component analysis (PCA) (Materials and Methods). Firing rates were partitioned into two non-overlapping segments: (1) from lever approach (2.5 s before press) to lever release, and (2) from lever release to reward consumption (1 s after nose poke). PCA subspaces were constructed separately for these periods. We projected population activity onto the top 4 PCs for visualization (Fig. 1m), revealing that individual PCs often aligned with distinct behavioral features. The top 4 PCs captured 71.1% and 75.7% of the variance for the press/release activity and retrieval activity, respectively. Press/release subspaces reflected the dynamics of lever approach and forepaw lifting (PCs 2, 3), holding (PCs 1–3), and release (PCs 1–4), while retrieval subspaces showed modulation peaking toward the nose poke (PCs 1–3). Based on the PC loading coefficients, we selected units with substantial contributions using *ad hoc* loading thresholds (Materials and Methods), which revealed average firing patterns tied to forepaw movements: lifting/lever press (L), holding (H), trigger/release (T), and reward retrieval movement (R) (Fig. 1n,o, fig. S2b,c).

There was a small overlap among these units (fig. S2d): 36.7% of retrieval units (33 out of 90) overlapped with forepaw-related units (fig. S2d). These results demonstrate that distinct DLS neuron subsets are recruited across various behavioral phases, covering forelimb movements for the SRT task and whole-body movement for reward retrieval.

Using chronic Neuropixels probes, we observed the sequential recruitment of DLS neurons, aligned with task events, in simultaneously recorded units during single sessions (Fig. 2a). In an example session, a total of 84 type-1 units (putative SPNs) and 18 other units were recorded. Of these, 64 type-1 units were modulated by behavioral events, the majority of which (57, 89.1%) were positively modulated. These units were modulated according to the task’s trial structure, with distinct behavior-relevant units intermixed along the dorsal-to-ventral axis of the striatum (Fig. 2a). The neurons exhibited repeated phasic activation across trials (Fig. 2b–d). After tracking the movement trajectories via DeepLabCut (*37*), we mapped spike rates onto the rat’s movement trajectories to reveal the movement dependence of these task-modulated spikes (Fig. 2e–g; see Materials and Methods, Equation 3). Some units (1 and 2) increased firing as the rat approached the lever, before lifting the forepaw; others fired during forelimb movements, such as reaching (unit 3), lever press (unit 4), or lever release (units 5 and 6).

**Fig. 2.**
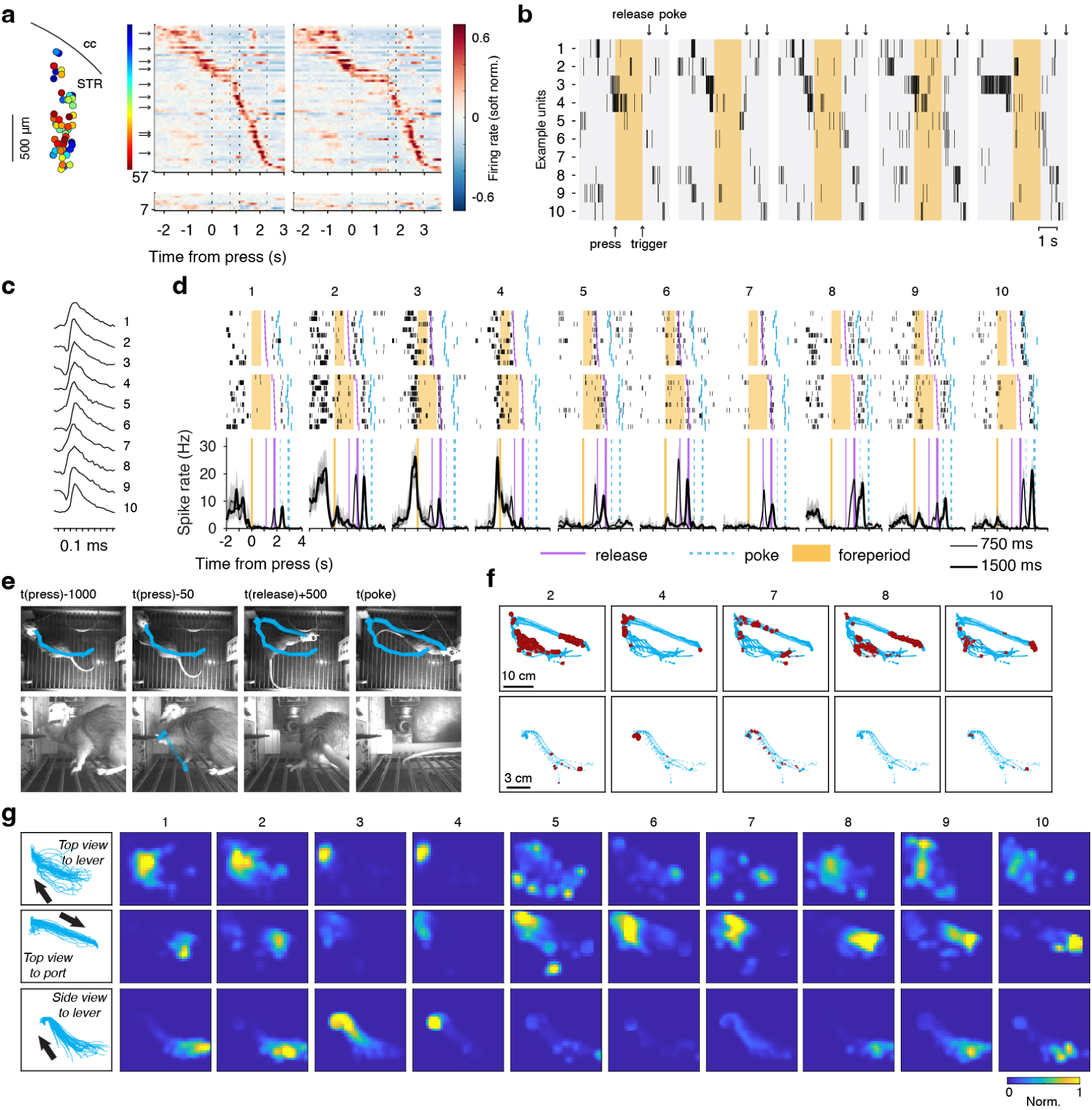
DLS neural activity tracks the rat’s movements during behavior. **a**, Neuropixels recordings from a single session. Units (putative SPNs) are arranged as in Fig. 1k. Arrows indicate 10 example units; dorso-ventral locations of 57 positively modulated units are color-coded by their response timings. The dorsal boundary of the striatum is shown (cc: corpus callosum; STR: striatum; relative to bregma: 0.68 mm). **b**, Spike rasters for 10 example units across five trials (FP = 1500 ms). Yellow rectangles denote the FP (press to trigger); arrows mark lever release and the port nose poke. **c–d**, Spike rasters and SDFs of the example units, as in Fig. 1i–j. **e**, Representative frames at four time points: 1000 ms and 50 ms before lever press, 500 ms after release, and port nose poke. Blue traces show head trajectories (first row, top view) and forepaw trajectories (second row, side view). **f**, Spikes (red) mapped onto trajectories for five example units. **g**, First column shows example movement trajectories. Other columns show spike rates mapped over space for 10 example units (each column is a unit). First row: head movement from lever approach to press; second row: head movement from trigger to port nose poke; third row: forelimb movement during lever reach and press. Spike rates are color-coded and normalized to *r*+*c*, where *r* is the 0.99 quantile of each unit’s spike-rate distribution and *c*= 1.

Additional neurons increased firing during reward retrieval (units 7–10), which involved turning the head toward the port. A similar pattern of activity was observed in another animal (fig. S3). Overall, DLS activity exhibited a task-dependent sequential structure aligned with specific forelimb and whole-body movements across discrete SRT phases, from lever approach to reward collection, consistent with the idea that task-related DLS spiking reflects, at least in part, low-level movement kinematics (*15*, *38*).

### Optogenetic inhibition of the striatum by activating interneurons

The striatum, like the cortex, contains local inhibitory interneurons that regulate the flow of information through its circuits (*39*, *40*). To probe the functional significance of task-related phasic activation of DLS neurons, we used an inhibitory interneuron-specific enhancer, mDlx, to introduce Channelrhodopsin-2 (ChR2) into striatal inhibitory interneurons (*41*, *42*) (Fig. 3a). Activating these interneurons temporarily inactivated SPNs with millisecond precision. The mDlx enhancer drives inhibitory interneuron-selective expression in the striatum with high fidelity (*42*). After injecting a recombinant adeno-associated virus (rAAV) encoding mDlx and ChR2 into the DLS, co-labeling of neurons expressing Foxp1 (an SPN-specific marker) and neurons expressing GFP was minimal: 25 out of 1813 GFP+ neurons (1.4%) expressed Foxp1, while 2 out of 3526 Foxp1+ neurons (< 0.1%) expressed GFP (data from two rats, 18 slices).

**Fig. 3.**
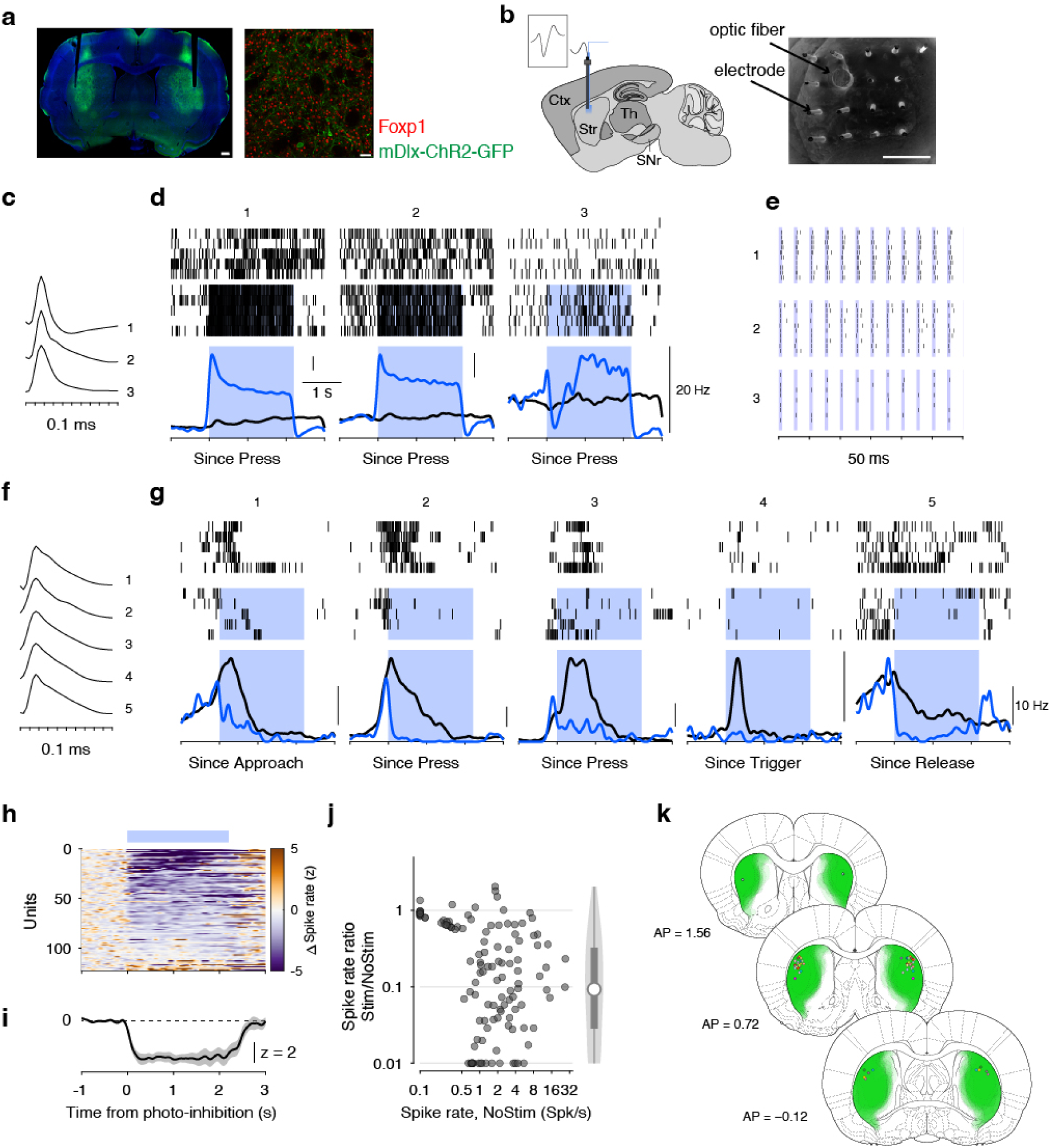
Optogenetic inhibition of the DLS via activation of inhibitory interneurons. **a**, Left: Brain slice illustrating viral expression of mDlx-ChR2-GFP in the DLS with an optical fiber track. Scale bar, 0.5 mm. Right: Foxp1 immunostaining (red) marks spiny projection neurons (SPNs), while green indicates ChR2-expressing inhibitory interneurons. Scale bar, 50 μm. **b**, Schematic of microwire array and optic fiber insertion in the DLS. Scale bar, 500 μm. **c**, Waveforms of three example ChR2+ units showing optogenetic activation, z-scored. **d**, Rasters for three ChR2+ units. Top: No photo-stimulation; middle: with photo-stimulation; bottom: spike density functions. Blue shading marks photo-stimulation periods (40 Hz, 5-ms pulse train, 2.2 s duration, coupled to lever press). **e**, Expanded view of stimulation, showing spikes tightly locked to 12 laser pulses (10 repeats shown). **f, g**, Same as **c**–**d** for five putative SPN units, demonstrating suppressed spiking during photo-stimulation in closed-loop conditions. **h**, Effect of the DLS photo-stimulation on 122 putative SPN units (wide waveforms), with z-scored differences in spiking between no-stimulation and photo-stimulation conditions. **i**, Average z-scored differences for the top 60 units from **h**. **j**, Ratio of spike rates in NoStim versus Stim conditions plotted against the spike rate in the NoStim condition. A violin plot with an embedded box plot is shown on the right (only units with baseline firing rate > 0.5 Hz are included). **k**, Fiber placements and viral expression across 26 rats.

To validate this approach physiologically, we performed *in vivo* optrode electrophysiological recordings from three rats implanted with 16-channel opto-microwire arrays (Fig. 3b). We performed electrophysiological recordings while rats engaged in the SRT task, and delivered laser pulse trains (40 Hz, 5-ms pulse width, 2.2-s duration) to the DLS at random task phases, such as during lever approach, in a small proportion of trials (< 25%). Putative inhibitory interneuron units were activated by laser pulses, showing elevated firing rates throughout the stimulation period (Fig. 3c–e). Units with wide-waveform spikes were suppressed during stimulation, and their behavior-related activation was reduced (Stim/NoStim spike rate ratio: median = 0.09, IQR: 0.03–0.32, *n*= 96 units with NoStim spike rate > 0.5 Hz; Fig. 3f–j). Taken together, DLS photo-inhibition via activation of DLS inhibitory interneurons produced sufficient inhibition of SPNs during behavior, enabling investigation of the effects of acute, behavior-coupled DLS inhibition.

Using this method, we combined closed-loop behavioral detection with optogenetics. For forelimb movements, we examined forelimb lift, lever press, lever holding, and lever release, key components of the lever-release SRT task (Figs. 4, 5, 6). For whole-body movements, we focused on the reward retrieval phase, spanning from lever release to nose poke (Fig. 7).

**Fig. 4.**
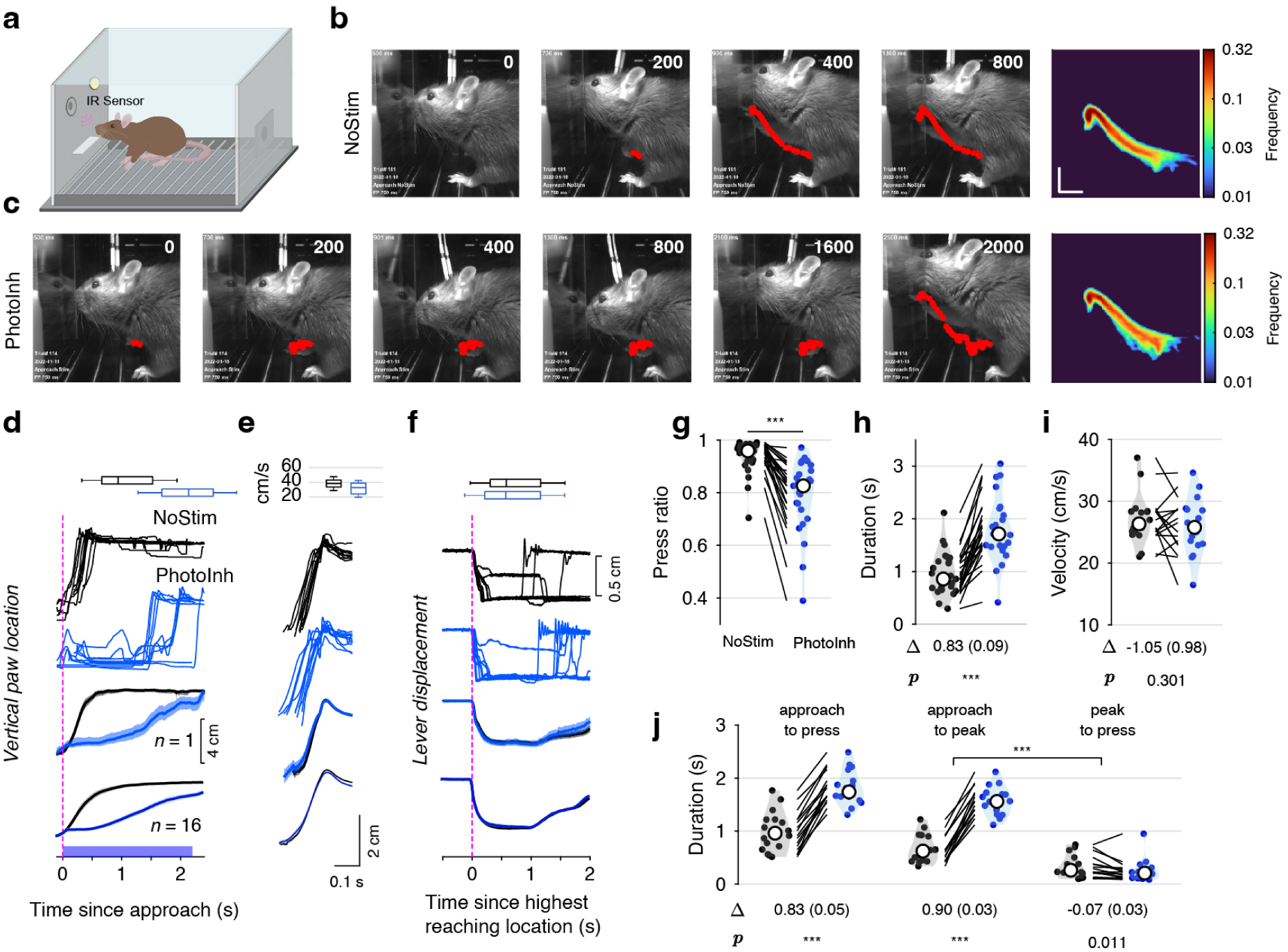
Bilateral DLS photo-inhibition delays the initiation of forelimb reaching. **a**, Schematic showing an infrared sensor detecting a rat’s approach to the lever. **b**, Sequential video frames showing a rat lifting its forepaw to press the lever. Frame numbers indicate time (ms) relative to approach detection (time = 0). Red dots trace the forepaw trajectory; the heat map (right) shows the across-trial frequency of forepaw crossings per grid square. Scale bar, 2.35 cm. **c**, Sequential video frames showing the forelimb trajectory during bilateral DLS photo-inhibition (starting at time 0, duration = 2.2 s). **d**, Top: horizontal box plots showing the distribution of approach-to-press durations across trials in an example rat (as shown in **b,c**). Rows 1–2: example forepaw vertical positions from the example rat (*n*= 1) for no-stimulation (NoStim; black) and DLS photo-inhibition (PhotoInh; blue) trials, aligned to approach detection. Row 3: average positions for the example rat (shading: 95% CI). Row 4: mean positions across 16 rats (shading: SEM). Blue rectangle denotes the laser stimulation period. **e**, As in **d**, but with forepaw vertical-position traces aligned to the time of the paw’s peak vertical position; box plots summarize forepaw velocity during the lifting phase. **f**, As in **d**, but showing lever-position traces aligned to the time of the paw’s peak vertical position; box plots summarize durations from peak to lever press. **g**, Ratio of successful lever presses within a 2.2-s window after approach detection (*n*=25 rats). *p*-value from a two-sided Wilcoxon signed-rank test. **h**, Time from approach detection to lever press (*n*= 25 rats). Values are estimated marginal means (EMMs, SEM in parentheses) for the PhotoInh−NoStim difference, derived from the LME model. **i**, Forepaw velocity during the lifting phase, as in **e** (*n*=16 rats). **j**, Time from approach detection to peak paw position and from peak position to lever press (*n*= 16 rats). Values are EMMs for PhotoInh − NoStim differences within each duration type, derived from an LME model, with *p*-values adjusted using Holm’s method. ∗∗∗ denotes *p*< 0.001.

**Fig. 5.**
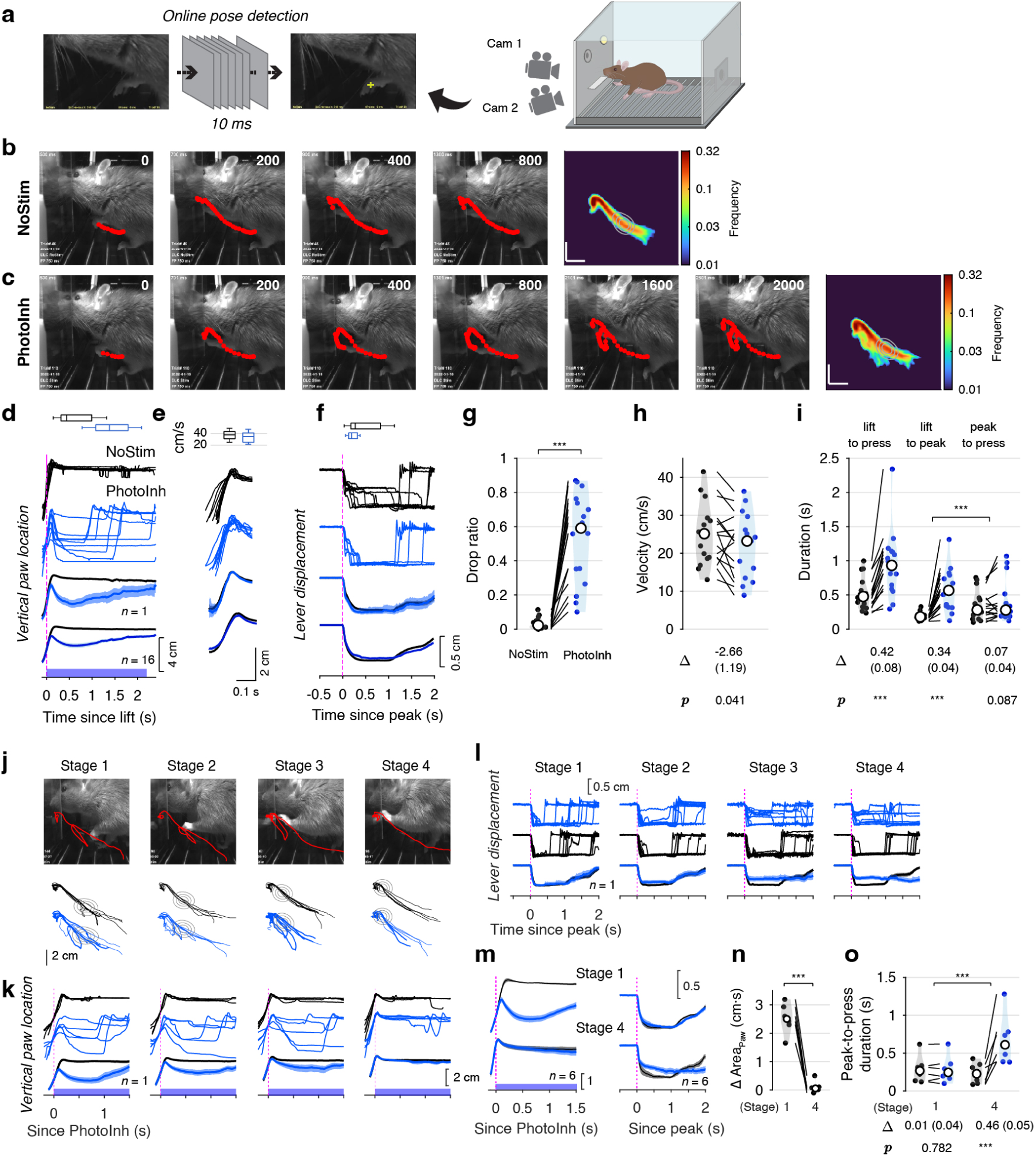
Bilateral DLS photo-inhibition interrupts the execution of forelimb reaching. **a**, Schematic showing online paw detection using a deep neural network. A dedicated camera captured frames at 50 fps over a small field of view; a second camera recorded high-speed video. **b**, Sequential video frames of a rat lifting its forepaw to press the lever. Frame numbers indicate time (ms) relative to lift detection (time = 0). Red dots trace the forepaw trajectory; the heat map (right) shows forepaw crossing frequency per grid square across trials. **c**, Sequential video frames showing the forepaw trajectory during bilateral DLS photo-inhibition (2.2-s duration, starting at time = 0). **d–f**, As in Fig. 4d–f, but with photo-inhibition applied at lift detection. **g**, Ratio of trials with paw dropping within a 2.2-s window after lift detection. Black dots, NoStim; blue dots, PhotoInh; white circle, median; lines connect paired data from the same rat (*n*= 16). **h**, Forepaw velocity during the lifting phase (*n*= 16 rats). **i**, Durations from lift detection to peak paw position, and from peak to lever press (*n*= 16 rats). **j**, Example frames showing four progressive stages of DLS photo-inhibition. Below each frame: example trajectories (black, NoStim; blue, PhotoInh). Gray contours represent isodensity lines of paw positions across trials at the onset of PhotoInh. **k**, Paw vertical positions for each PhotoInh stage in the example rat from **j**. Time = 0 marks PhotoInh onset. **l**, Lever positions for each PhotoInh stage in the example rat, aligned to the time of the paw’s peak vertical position. **m**, Averaged paw vertical positions and lever positions from six rats across two PhotoInh stages. **n**, Integrated differences in paw vertical position across the two PhotoInh stages. *P*-value from paired *t*-test. **o**, Peak-to-press durations across the two PhotoInh stages.

**Fig. 6.**
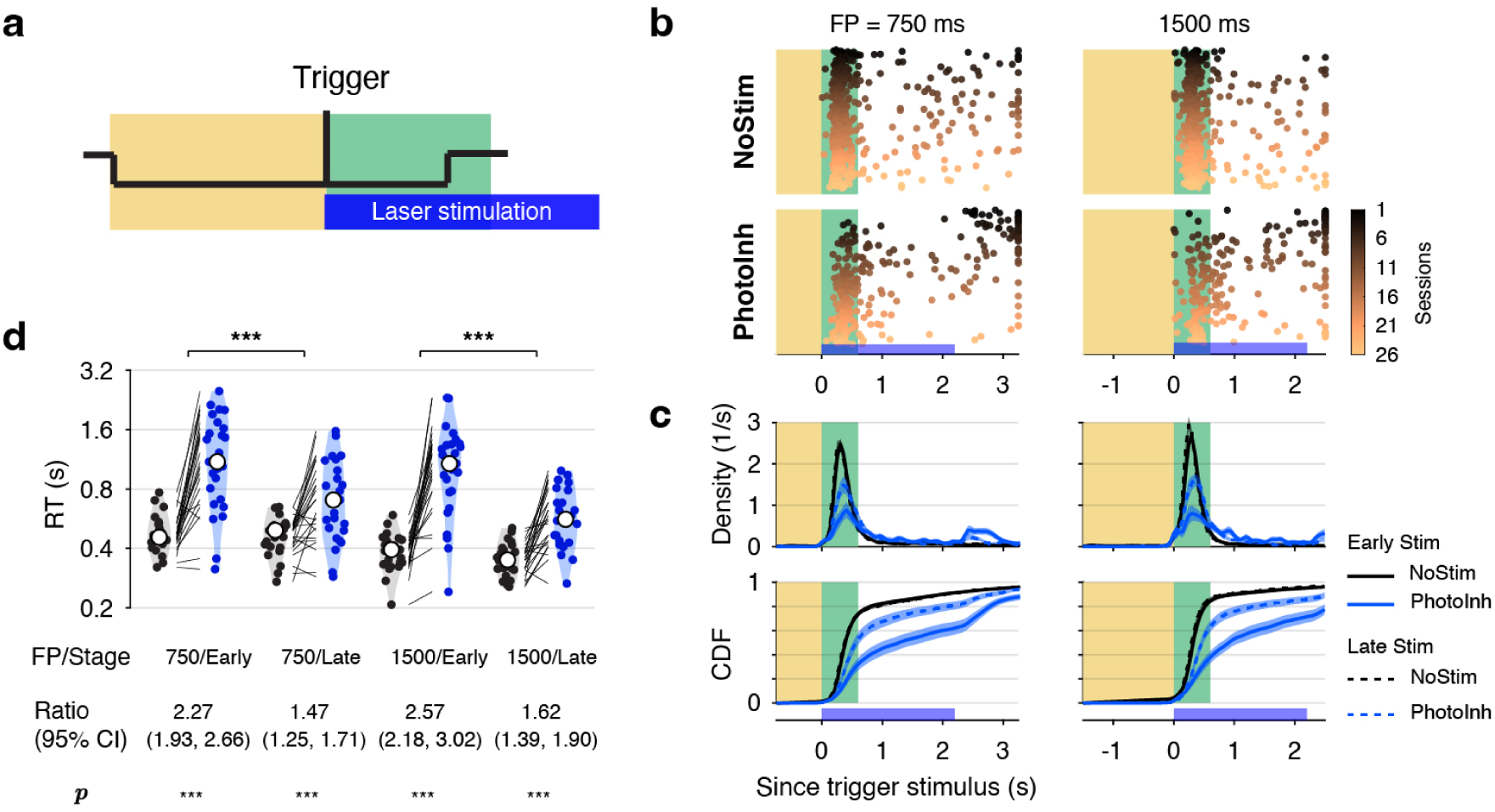
DLS photo-inhibition prolongs reaction time. **a**, Schematic of bilateral DLS photo-inhibition coupled to the trigger stimulus, applied only if no premature response occurred during the FP. **b**, Lever-release responses in an example rat with (blue) or without (gray; 20% of trials plotted) bilateral DLS photo-inhibition, color-coded by session. Left: short FP (750 ms); right: long FP (1500 ms). **c**, Empirical probability density functions (PDFs) and cumulative distribution functions (CDFs) of reaction times for *n*= 25 rats. The blue rectangle marks the laser stimulation period. Solid lines: early photo-inhibition (≤ 60 trials); dashed lines: late photo-inhibition (> 60 trials); shading denotes SEM. **d**, Geometric mean reaction times with (blue) or without (gray) trigger-coupled photo-inhibition from *n*= 25 rats, grouped by FP and photo-inhibition history (Early, Late). Violin plots show distributions; white circle, median; RT ratios (PhotoInh/NoStim), 95% confidence intervals (CI), and Holm-adjusted *p*-values from pairwise EMM contrasts in an LME model are shown below.

**Fig. 7.**
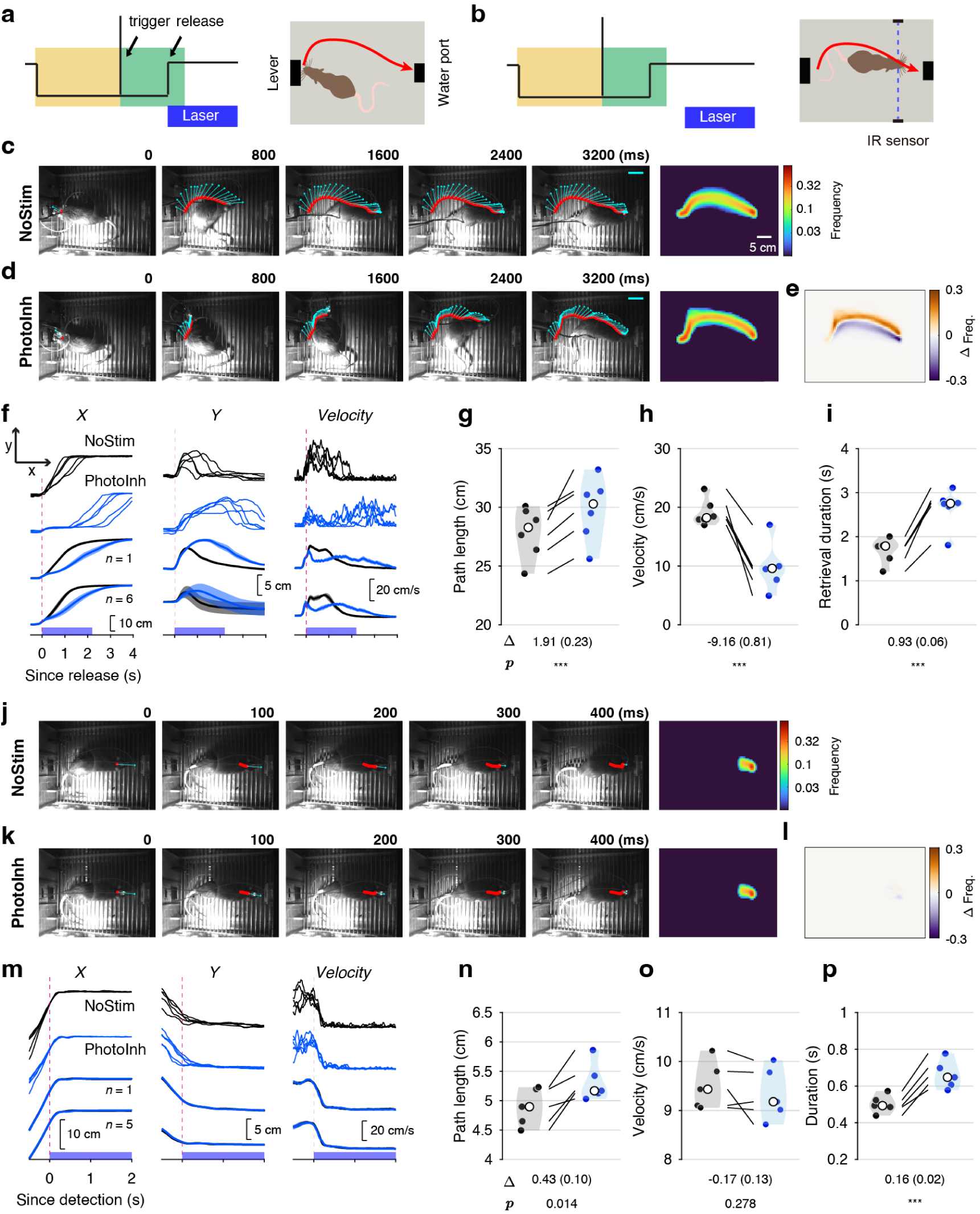
DLS photo-inhibition impairs whole-body movements during reward retrieval. **a**, Left: bilateral DLS photo-inhibition coupled to lever release. Right: Top-view camera tracks locomotion from lever release to port nose poke. **b**, Left: bilateral DLS photo-inhibition coupled to an infrared sensor detecting the rat’s approach to the water port. **c**, Sequential video frames at time points (ms) after lever release in a trial without photo-inhibition from a representative rat. Cyan arrows indicate head direction (50-ms intervals) with arrow length denoting movement speed (scale bar, 20 cm/s); red dots mark head position. Right: trajectory heat map (color indicates pixel traversal frequency). **d**, As in **c**, but for a bilateral photo-inhibition trial (stimulation onset at time = 0). **e**, Difference in trajectory heat maps (**d** minus **c**). **f**, Movement along the X-axis (toward the water port), Y-axis, and instantaneous velocity. Rows 1–2: single trials from the representative rat. Row 3: averages with 95% confidence intervals (shaded) for the representative rat (*n*= 1). Row 4: averages with SEM (shaded) across *n*= 6 rats. The blue rectangle marks the laser stimulation period. **g–i**, Movement path length, mean velocity, and retrieval duration for six rats (dots: individual-rat averages; white circle: group medians). Mean differences (SEM in parentheses) and Holm-adjusted *p*-values are from pairwise EMM contrasts based on an LME model. **j–p**, As **c–i**, but for bilateral photo-inhibition near the water port, as shown in **b**.

### DLS inhibition delays forelimb reaching

To examine how DLS inhibition affects the initiation and execution of a lever press, a forelimb movement, we installed an infrared sensor above the lever. Once triggered, the sensor initiated delivery of a 2.2-s, 40-Hz pulse train (5-ms pulses) to the DLS in a small proportion (10%) of trials (Fig. 4a). As shown in the example, bilateral DLS photo-inhibition (PhotoInh) delayed movement initiation compared to uninhibited trials (NoStim) (Fig. 4b–d; see Movie S1). The press ratio, defined as the proportion of trials in which the rat pressed the lever within a 2.2-s window after approach detection, decreased by 0.141 (median paired difference; 95% CI: [0.1, 0.185]; *p* < 0.001, two-sided Wilcoxon signed-rank test, *n*= 25 rats; Fig. 4g).

We measured the time from approach detection to lever press (*t*_a2p_) and fitted a linear mixed-effects (LME) model (Equation 4) with stimulation type (StimType: NoStim or PhotoInh) as a fixed effect. The model included random intercepts and random slopes for StimType by rat to account for individual variability. Bilateral DLS photo-inhibition significantly increased *t*_a2p_ (β_1_ = 0.83±0.09 s, mean ± SEM, *t*(24) = 9.42, *p* < 0.001, *n*= 25 rats; Fig. 4h).

When successfully initiated, lever-reaching trajectories during PhotoInh resembled those in NoStim trials (Fig. 4b,c,e). To assess forepaw movement velocity, in a subset of rats with unobstructed forepaw tracking (*n*= 16), we fitted an LME model including StimType as a fixed effect (Equation 5). The estimated marginal mean velocity was 27.07 ±1.02 cm/s for NoStim (mean ± SEM), comparable to 26.02±1.15 cm/s for PhotoInh. The coefficient (β_1_) for StimType was −1.05±0.98 cm/s (*t*(14) =−1.07, *p* = 0.301), suggesting a small and inconsistent population-level effect of DLS photo-inhibition on forepaw velocity.

For each trial, we partitioned *t*_a2p_ into two durations: from approach to lift peak (*t*_a2pk_) and from lift peak to lever press (*t*_pk2p_). We then fitted an LME model to assess the interaction between the effect of photo-inhibition (StimType) and duration type (DurationType; Equation 6). A strong StimType × DurationType interaction was observed (β_3_ =−0.97 ±0.02 s, *t*(8127) =−51.81, *p* < 0.001, *n*= 16 rats), indicating that photo-inhibition affected these two durations differently. The increase in *t*_a2p_ (0.83±0.05 s) was entirely attributable to the increase in *t*_a2pk_ (0.90±0.03 s), but not the subsequent *t*_pk2p_ (−0.07 ±0.03 s; Fig. 4j). Lever movement was monitored with a piezo sensor, and lever position was reconstructed (Fig. 4f). Lever movements were indistinguishable between NoStim and PhotoInh trials (Fig. 4f), suggesting that a similar force was applied to the lever. Together, DLS photo-inhibition delayed forepaw lifting. However, once initiated, forepaw reaching and subsequent lever press showed similar kinematics and vigor (velocity and force) under both NoStim and PhotoInh conditions.

The results above were obtained with bilateral DLS photo-inhibition. In 10 rats, we tested unilateral photo-inhibition (contralateral or ipsilateral to the lever-pressing paw) and compared its effects on *t*_a2p_ with those of bilateral photo-inhibition (Equation 7, fig. S4a–e). Unilateral photo-inhibition also significantly decreased the press ratio and increased *t*_a2p_ relative to NoStim. Overall, bilateral photo-inhibition had the strongest effect, followed by contralateral and then ipsilateral inhibition. These findings support a role for the DLS in initiating forelimb actions, suggesting that phasic pre-lift DLS activity contributes causally to movement initiation (e.g., example units 1–2 in Fig. 2).

### DLS inhibition interrupts execution of forelimb reaching

From the perspective of action selection (*12*, *14*), forelimb lever reaching can be viewed as a single action chosen among alternatives, such as standing still, turning away, or using the contralateral paw. Once a lever-reaching movement has been initiated, we asked whether DLS photo-inhibition modulates its ongoing execution. Recent work suggests that the striatum not only selects actions but also encodes, and is critical for producing, complex forelimb kinematics (*15*, *16*). However, the causal contribution of DLS activity during a single reach remains largely untested.

To achieve this, we implemented real-time, high-speed video-based pose detection with online estimation computed within 10 ms per frame at 50 Hz (Fig. 5a) (*43*, *44*). In a subset (10%) of trials, paw lift triggered bilateral DLS photo-inhibition. Lift-triggered and approach-triggered photo-inhibition trials were often interleaved within the same session.

As shown in the example, DLS photo-inhibition at lift onset interrupted reach execution, causing the paw to drop and hover or pause for variable durations (Fig. 5b–d; see Movie S2). The paw-drop ratios increased by 0.499 (median paired difference; 95% CI: [0.353, 0.683]; *p* < 0.001, two-sided Wilcoxon signed-rank test, *n*= 16 rats; Fig. 5g). Rats often re-initiated reaching during PhotoInh (Fig. 5c–e), but with a slight velocity reduction (10.6±5.9% decrease; Fig. 5h), likely because of mid-trajectory restarts. Despite these interruptions, once reaching resumed, subsequent lever pressing was largely unimpaired (Fig. 5f).

As above, we partitioned the lift-to-press duration (*t*_l2p_) into the lift-to-lift-peak (*t*_l2pk_) and lift-peak-to-press (*t*_pk2p_) segments, and fitted an LME model (Equation 9). The model revealed a significant StimType × DurationType interaction (β_3_ =−0.27 ±0.02 s, mean ± SEM, *t*(5444) =−12.55, *p* < 0.001), indicating that photo-inhibition affected the two durations differently. Photo-inhibition increased *t*_l2p_ (0.42±0.08 s; Equation 8) primarily because of a delay in *t*_l2pk_ (0.34±0.04 s), with only a small effect on *t*_pk2p_ (0.07 ±0.04 s) (Fig. 5i).

We next designed an experiment to test the effect of photo-inhibition at different points along the reaching movement (Fig. 5j). As photo-inhibition onset was progressively delayed in the reaching movement, the paw-drop rate decreased (Fig. 5j–k), but lever pressing became more impaired (Fig. 5l). In six rats, early-stage photo-inhibition altered forelimb trajectories (Fig. 5m left, n), whereas late-stage photo-inhibition more strongly affected lever movements (Fig. 5m right). LME modeling of *t*_pk2p_ (Equation 10) revealed a StimType × Stage interaction (β_3_ = 0.45± 0.04 s, mean ± SEM, *t*(967) = 10.7, *p* < 0.001), confirming a larger effect on *t*_pk2p_ at a later stage (0.46±0.05 s) than at the early stage (0.01±0.04 s) (Fig. 5o). Together, these findings indicate that DLS modulates movement execution in a time-sensitive manner: the closer inhibition occurs to movement onset (immediately before or after), the greater the impairment in both initiation and execution.

The results above were based on bilateral DLS photo-inhibition. Compared with bilateral photo-inhibition, unilateral photo-inhibition had weaker effects on the paw-drop rate and on *t*_l2p_ (Equation 11, fig. S4f–j). Relative to the lifting paw, contralateral photo-inhibition had a larger effect than ipsilateral photo-inhibition.

### DLS inhibition prolongs reaction time

A prominent component of DLS neural dynamics correlated with lever-release responses following the trigger stimulus (e.g., trigger-release units in Fig. 1n and fig. S2b). To assess its functional role, we suppressed this activity by coupling laser onset to the trigger stimulus in rats (Fig. 6a) and examined the effects of DLS photo-inhibition on reaction time (lever release; see Movie S3) and on the subsequent reward retrieval process (see below).

Trigger-coupled bilateral DLS photo-inhibition prolonged reaction times (β_1_ = 0.69±0.1, mean ± SEM, *t*(63.5) = 7.25, *p* < 0.001, *n*= 25 rats; LME model in Equation 12), initially by 2.27-fold (750 ms FP) and 2.57-fold (1500 ms FP) (Fig. 6d; Early Stim). This slowing effect attenuated with repeated inhibitions, reducing to 1.47-fold and 1.62-fold (Fig. 6d; Late Stim), suggesting partial adaptation. Compared to bilateral DLS photo-inhibition, unilateral photo-inhibition had inconsistent effects (fig. S5). Overall, DLS activity coupled to trigger-release is crucial for initiating the forelimb lever-release responses.

### DLS inhibition reduces locomotor efficiency and motor vigor during reward retrieval

In the behavioral box, after a correct lever release, rats retrieved water rewards from a reward port on the wall opposite the lever, requiring body turning, locomotion, and a self-initiated nose poke (Fig. 1a,d). Reward retrieval duration was defined as the time between lever release and nose poke at the reward port. Retrieval duration did not affect reward size. To assess the impact of DLS photo-inhibition on reward retrieval, we coupled the onset of photo-inhibition to lever release (Fig. 7a) and tracked head coordinates and direction using a top-view camera (Fig. 7a).

Bilateral DLS photo-inhibition prolonged the reward retrieval durations, with trajectory frequency maps showing longer, more outward paths and reduced movement velocity (Fig. 7d–f; see y position and velocity in Fig. 7f; see Movie S4). Using an LME model with fixed effects including StimType (NoStim, PhotoInh) and StimSide (bilateral, ipsilateral, contralateral to turning direction) and random effects for animals (*n*= 6; Equation 13), we found that bilateral photo-inhibition significantly increased path length (β_1_=1.91±0.23 cm, *t*(7.5)=8.34, *p*<0.001), reduced velocity (β_1_ =−9.16±0.81 cm/s, *t*(6.6) =−11.3, *p* < 0.001), and prolonged reward retrieval duration (β_1_ = 0.93±0.06 s, *t*(7.4) = 15.28, *p* < 0.001), indicating impaired movement efficiency (larger outward turns) and reduced movement vigor (slower velocity). Compared to bilateral inhibition, unilateral inhibition had similar but weaker effects, with ipsilateral inhibition producing a larger effect than contralateral inhibition (fig. S6).

When DLS photo-inhibition was instead coupled to the trigger stimulus (Fig. 6), rats, despite longer reaction times, sometimes released the lever within the response window, allowing us to test whether the subsequent reward retrieval process was affected (fig. S7; same example rat as in Fig. 7c). The effects were similar: trigger-coupled DLS photo-inhibition also increased the reward retrieval duration, particularly with bilateral inhibition (β_1_=0.52±0.04 s, *t*(24.6)=12.42, *p*<0.001, *n*=25 rats; fig. S7b,c). Movement tracking in a subset of rats (*n*=11; fig. S7d–j) showed increased path length and reduced velocity, similar to Fig. 7c–i (see Movie S5).

Finally, using an infrared sensor near the reward port, we applied bilateral photo-inhibition toward the end of retrieval locomotion but before the nose poke (*n*= 5 rats; Fig. 7b; see Movie S6). This occurred as rats began to decelerate (see velocity in Fig. 7m) and prepared to initiate the nose poke. Photo-inhibition had little effect on the remaining trajectory (Fig. 7j–m), only slightly increasing the path length leading to the nose poke (β_1_ = 0.43±0.1 cm, *t*(4) = 4.17, *p* = 0.014, Fig. 7n). Photo-inhibition did not significantly change the movement velocity (β_1_ =−0.17 ±0.13 cm/s, *t*(3.6) =−1.28, *p* = 0.278), suggesting normal deceleration. The duration from detection to nose poke onset increased slightly (β_1_ = 0.16±0.02 s, *t*(4.6) = 7.68, *p* < 0.001, Fig. 7o–p). Overall, DLS photo-inhibition subtly delayed nose poke behavior, in contrast to its much stronger effects on forelimb lifting during lever pressing (Figs. 4 and 5). Thus, nose-poke-related striatal spikes, despite their abundance (e.g., PC2 in the reward/poke subspaces was almost entirely driven by nose poke near the reward port; Fig. 1m), appear to contribute only mildly to driving the nose poke movement itself.

### Control experiments

To minimize the visual confounds from laser stimulation, as in previous optogenetics studies (*45*, *46*), all optogenetic experiments in this study were conducted with a masking blue-LED flash (20 Hz, 4–6 s) mounted on the ceiling of the box, applied during both NoStim and PhotoInh trials and beginning at lever approach. Control experiments were performed in rats (*n*= 4) injected with an AAV expressing GFP but lacking ChR2. These rats underwent identical training, behavioral testing, and analysis (fig. S8). Briefly, in control rats, DLS photo-stimulation (i.e., sham photo-inhibition), when coupled to lever approach, did not alter the approach-to-press duration (fig. S8a–e). However, when coupled with the trigger tone, it slightly shortened reaction time (fig. S8f–g), likely because of the additional visual cue from the laser coinciding with the tone. DLS photo-stimulation had no effect on whole-body locomotion toward the reward port (fig. S8h–n). Thus, the observed behavioral effects of DLS photo-inhibition were attributable to modulation of DLS and downstream activity, rather than to nonspecific factors such as tissue heating (*47*).

### DMS-DLS dissociation

In addition, we conducted parallel experiments and analyses in the dorsomedial striatum (DMS) using the same methodology and pipeline (*n*= 12 rats; Fig. 8a-b). DMS photo-inhibition produced no detectable change in approach-to-press duration. That is, forelimb movement initiation was unimpaired (Fig. 8c–i). Lift-coupled photo-inhibition was not performed in DMS.

**Fig. 8.**
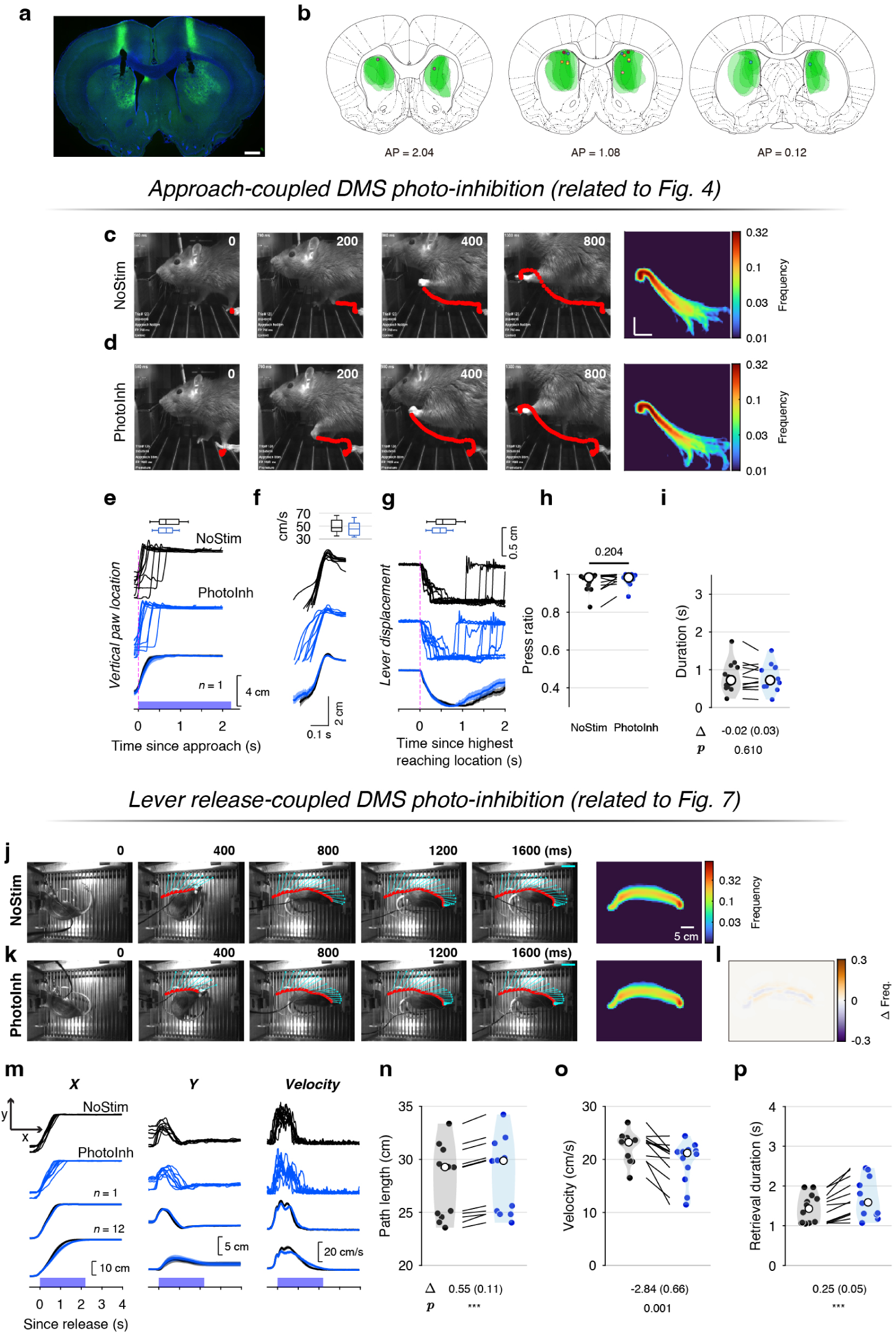
Minimal effects of bilateral DMS photo-inhibition during approach and reward retrieval. **a**, Brain slice illustrating viral expression of mDlx-ChR2-GFP in the DMS with an optical fiber track. Scale bar, 1 mm. **b**, Fiber placements and viral expression across 8 rats. **c–i**, As in Fig. 4b–h but for rats with DMS photo-inhibition (*n*= 12) with mDlx-ChR2-GFP expressed in inhibitory interneurons of the DMS. In these experiments, DMS photo-inhibition was coupled to lever approach, and statistical analyses were conducted as in the DLS photo-inhibition experiments. **j–p**, As in Fig. 7c–i, with DMS photo-inhibition coupled to lever release.

When photo-inhibition was coupled to lever release, whole-body movement velocity slowed and reward retrieval duration increased, but these effects were substantially smaller than with DLS photo-inhibition (Fig. 8j–p). Taken together, both the lateral (DLS) and medial (DMS) domains of the dorsal striatum contribute to whole-body movement, whereas the DLS additionally gates and controls forelimb reach initiation and execution.

### DLS inhibition results in mixed responses in BG output (SNr)

The behavioral effects of striatal photo-inhibition ultimately manifest as changes in BG output via structures such as the SNr (*48*, *49*), with downstream effects on targets including the motor thalamus, superior colliculus, and brainstem (*50–54*). Increased BG output inhibits downstream circuits, delaying forelimb movement initiation (*50*) or suppressing repetitive licking (*55*). Decreasing BG output slows reaching movements in monkeys (*24*). Thus, perturbing BG output in either direction can impair movement.

To examine the effects of DLS inhibition on BG output, we performed tetrode recordings from the SNr of nine rats during the SRT task. In six of these rats, we bilaterally implanted optical fibers in the DLS to enable simultaneous SNr recordings during DLS photo-inhibition (Fig. 9a). Spike waveforms recorded in the SNr exhibited a continuum of widths and shapes, which we roughly grouped into two types (1 and 2) based on spike width (type 1: 0.1±0.02 ms, *n*= 249; type 2: 0.18±0.03 ms, *n*= 64; mean ± SD) and their projections onto the first principal component (Figure 9b,c). These unit types were spatially intermixed across recording depths and exhibited similar baseline firing rates (Figure 9d,e), possibly corresponding to genetically defined cell types such as parvalbumin (PV)-positive and non-PV neurons (*51*, *56*) or those projecting to specific targets (*52*). Using the same criteria applied to DLS units, including ISI violations and event modulation, we selected 240 (type 1) and 60 (type 2) SNr units for further analysis.

**Fig. 9.**
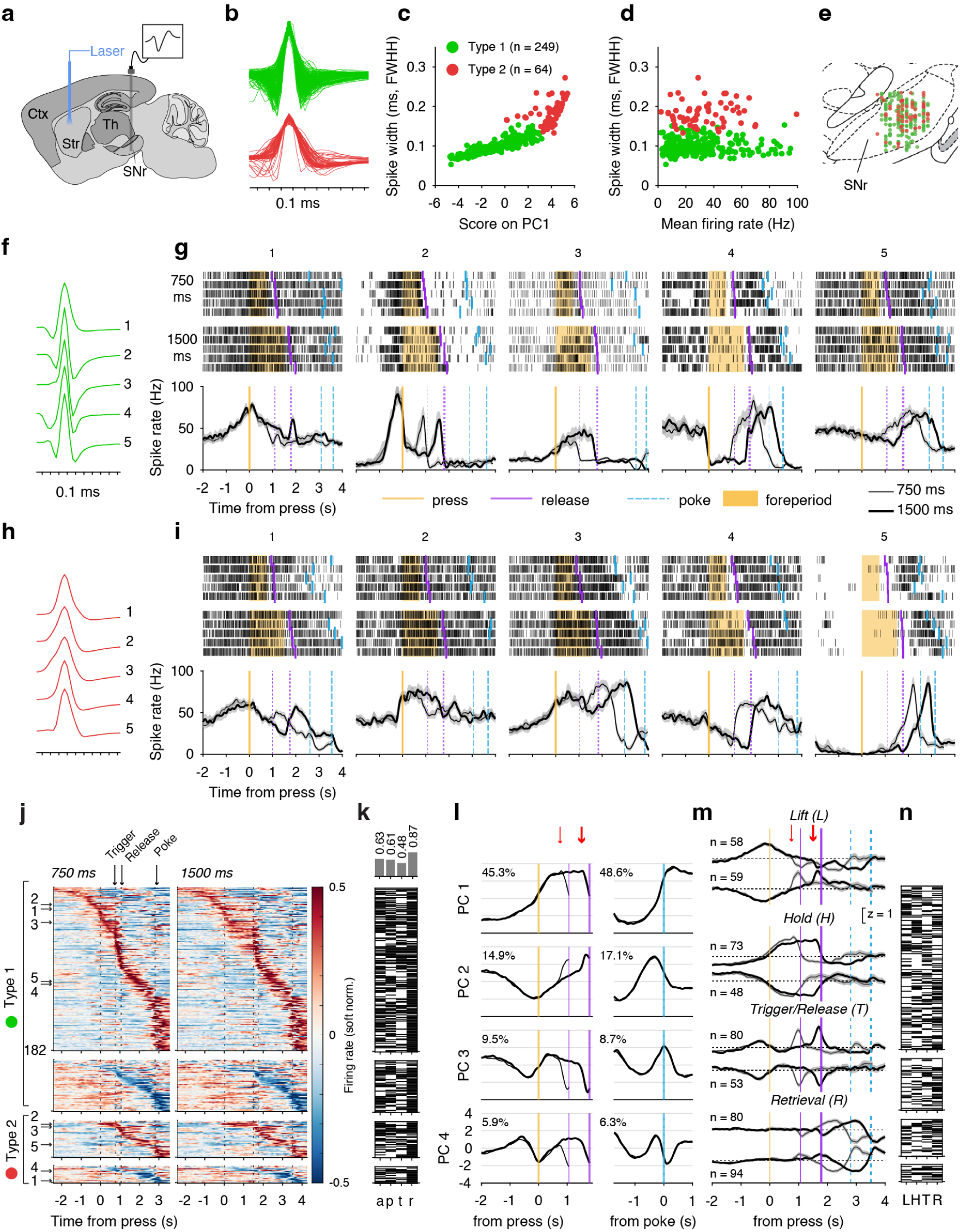
Neural activity in the substantia nigra pars reticulata (SNr) during the SRT task. **a**, Schematic of tetrode recordings in the SNr, often combined with striatal photo-inhibition. **b**, Two waveform types (1 and 2) recorded in the SNr with different widths; waveforms are plotted with negative voltage upward. **c**, Spike width (FWHM) plotted against the projection onto the first principal component. **d**, Spike width (FWHM) plotted against average firing rate. **e**, Estimated anatomical locations of recorded units in the SNr. **f**, Example waveforms from five type-1 units. **g**, Spike rasters and SDFs for the example units in **f**. Same layout as in Fig. 1i. **h–i**, Same as in **f–g**, but for five example type-2 units. **j**, SDFs (soft-normalized) of SNr units: rows 1–2 show positively and negatively modulated type-1 units; rows 3–4 show type-2 units. **k**, Modulation by event for each unit based on GLM fits (FDR= 0.05). Unit order is the same as in **j**. The histogram shows the proportion of units modulated by each event. **l**, Population projections onto the first four PCs around lever press (left) or port nose poke (right). Thin/thick lines represent 750/1500 ms FP. Colors match those in **j**; numbers indicate variance explained. Red arrows indicate trigger onset. **m**, Average z-scored SDFs of functional units selected based on their projection loadings (see Materials and Methods). **n**, Index of selected functional units (Lift, Hold, Trigger/Release, or Retrieval); horizontal lines indicate unit assignments, ordered from top to bottom to match the colormap order in **j**. Shading denotes SEM in **l** and **m**.

Consistent with earlier studies (*25*, *50*, *57*, *58*), SNr neurons exhibited heterogeneous firing patterns during behavior, with behavior-relevant increases or decreases in firing rate from a high baseline, modulated by events such as lever approach, press, holding, release, and reward retrieval (Figure 9f–k). Both type-1 (narrow waveform) and type-2 (broad waveform) SNr neurons were modulated. PCA identified subspaces capturing variance in SNr population activity modulated by forelimb responses or reward retrieval, with population projections similar to those observed in the DLS (Fig. 9l). As in Figure 1n, we used these PCA subspaces to guide unit selection based on their loadings (Fig. 9m, n). Notably, SNr units exhibited more mixed selectivity (Figure 9k,n; fig. S2g): 85.6% of retrieval units overlapped with lift/hold/trigger units, a higher proportion than observed in the DLS units (36.7%, fig. S2d; χ^2^(1, 1) = 61.2, *p* < 0.001, chi-square test). These findings suggest convergence and mixing of behaviorally relevant modulation from the DLS onto SNr neurons.

In a subset of SNr recordings (*n*= 124 units), press-aligned DLS photo-inhibition was applied in a small fraction of trials. We aligned activity to lever press because, although premature responses increased with DLS photo-inhibition, they typically emerged only after repeated stimulation and increased with longer holding durations, rather than immediately after the press (fig. S9). We selected trials in which the holding duration was at least 500 ms for both NoStim and PhotoInh conditions, ensuring that the first 500 ms of behavior was matched across conditions to allow for direct comparison of neural activity (Figure 10a–d). As shown in these examples, DLS photo-inhibition induced either an increase or a decrease in the firing rate of single SNr neurons (Fig. 10e–f), and the distribution of change directions did not differ significantly between type 1 and 2 neurons (χ^2^(2) = 0.22, *p* = 0.895, chi-square test; Fig. 10g).

**Fig. 10.**
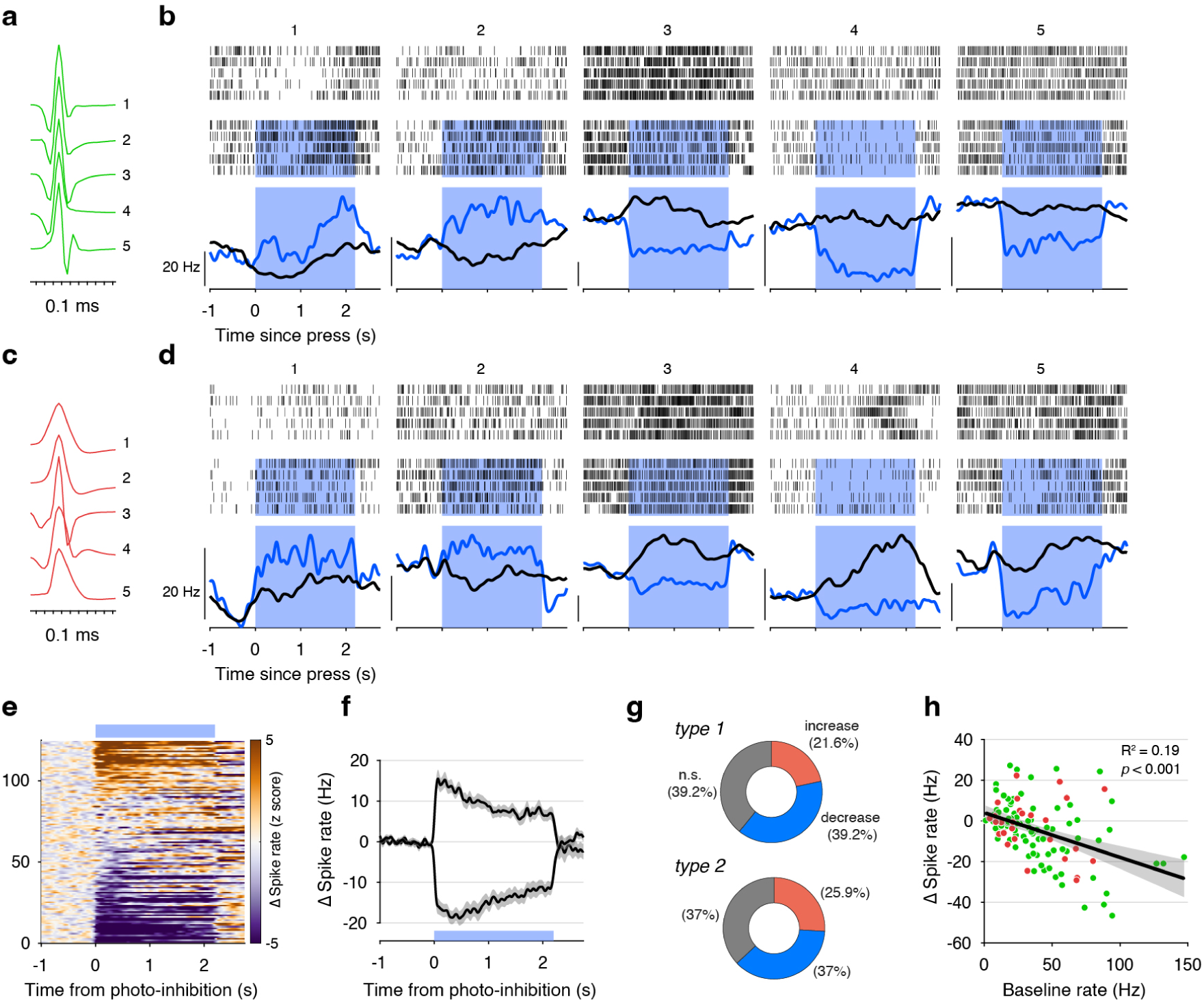
DLS photo-inhibition bidirectionally modulates spike rates in SNr neurons. **a**, Example waveforms from five type-1 SNr neurons. **b**, Spike rasters and SDFs (black, NoStim; blue, PhotoInh) for five example type-1 SNr neurons with and without DLS photo-inhibition, aligned to lever-press onset. The blue rectangles mark the laser stimulation periods. **c–d**, Same as **a–b** for five type-2 SNr neurons with wider waveforms. **e**, Z-scored SDF differences for 124 neurons(PhotoInh −NoStim), ordered by firing-rate changes during photo-inhibition, from increased firing (top) to decreased firing (bottom). **f**, Mean SDFs for 28 neurons with increased firing rates versus 48 neurons with decreased firing rates during DLS photo-inhibition. **g**, Distribution of type-1 and type-2 neurons exhibiting increased or decreased firing rates during DLS photo-inhibition. **h**, A negative correlation between the baseline (NoStim) firing rate and the change in firing rate induced by DLS photo-inhibition.

The change occurred approximately 14 ms (time at half maximum) for neurons with decreasing firing rates and 8 ms for neurons with increasing firing rates (Fig. 10f). The complex and diverse effects of DLS inhibition on SNr neurons observed here are consistent with previous studies involving striatal activation or inactivation (*48*, *49*).

There was a significant negative correlation between the firing rate (FR) in the NoStim condition and the modulation magnitude (FR_PhotoInh_ −FR_NoStim_) under photo-inhibition (Spearman’s ρ = −0.43, df = 122, *p* < 0.001), indicating that neurons with higher baseline firing rates tended to decrease their activity during DLS photo-inhibition. By contrast, neurons with lower baseline firing increased their activity (Fig. 10h).

## Discussion

Our experiments revealed a continuous involvement of DLS neural activity in the expression of learned forelimb and whole-body movements. DLS inhibition delayed the initiation of forelimb movements, including reaching and lever release, and disrupted their ongoing execution.

Whole-body locomotion was slowed throughout the inhibition period and followed less efficient trajectories toward the reward port. The nose-poke movement, by contrast, appeared less dependent on DLS activity. Together, our results demonstrate that DLS spiking causally contributes to movement initiation and execution, while also revealing conditions under which robust DLS activity, although present, is not critical for generating specific movements.

### Neural activity in DLS and SNr is linked to diverse movement types

In our task, the lever and the reward port were placed on opposite sides of the behavioral box (32 or 35 cm apart), naturally segregating lever-related forelimb movements from reward-oriented whole-body locomotion. Individual DLS neurons were selectively coupled to one or more discrete behavioral events such as lever approach, paw lift (reach), lever press and hold, lever release, and reward retrieval. Neural activity in the BG output nucleus, the SNr, was also modulated by these events. Comparing the activity patterns, using either event-modulation or PCA-based unit selection revealed more mixed selectivity in the SNr (for example, compare Fig. 1l,o and Fig. 9k,n). That is, a single SNr neuron is more likely to be modulated at multiple task events and to be linked to several movement types.

Photo-inhibition of DLS neurons produced mixed effects in the SNr: some neurons increased their firing rate, while others decreased it, consistent with previous work (*48*, *49*). The firing rate of an SNr neuron is determined by many factors, including its intrinsic firing properties independent of synaptic inputs (*52*), inhibitory input from the direct and indirect pathways, and excitatory input from the subthalamic nucleus (STN)(*59*, *60*). Although the precise rules of connectivity from the striatum to the SNr remain largely unexplored, our results suggest that information from multiple striatal inputs converges onto single SNr neurons, producing multiplexed firing patterns. Withdrawal of DLS input disrupted the normal firing patterns of SNr neurons, which impaired movement. Indeed, abnormal output patterns in the BG may underlie striatum-related disorders such as Parkinson’s disease (*61*).

### Continuous control of forelimb and locomotor movements by DLS

DLS photo-inhibition produced differential effects on forelimb and whole-body locomotor movements. When applied before movement onset, DLS photo-inhibition delayed forelimb reaching or tone-cued lever release, consistent with a role in action selection and initiation (*14*, *62*). After reach initiation, DLS photo-inhibition interrupted ongoing execution, indicating that the DLS also contributes to the progression of an initiated movement. Notably, in both cases, reaches were often re-initiated during inhibition and completed at similar velocities and along similar trajectories as in uninhibited trials. In comparison, whole-body locomotion did not stop completely during photo-inhibition: rats continued toward the reward port at lower speeds and via larger circles, indicating reduced movement vigor (defined as the “speed, amplitude, or frequency” of a movement (*17*)). A possible interpretation is that the alternating movements of the forelimbs and hindlimbs during whole-body locomotion, particularly during turning, were repeatedly disrupted during DLS inhibition, albeit to a lesser extent compared to forelimb reaching. The reduced movement speed may thus reflect these impairments in movement initiation and execution.

Finally, not all striatal spikes are involved in driving a movement. In our task, nose-poking into the reward port, the final step of a behavioral trial, was represented by a substantial fraction of DLS spikes; however, the movement was largely preserved during DLS photo-inhibition: animals could perform the nose-poke movement with little deficit in initiation or vigor during DLS inhibition. The function of these spikes is thus not to drive the movement itself. Previously, it has been reported that activity associated with the final step of a behavioral trial, often also coupled to reward retrieval, increases in proportion with learning (*9*). Although the emergence of such activity has been associated with habit formation or skill acquisition, its function has not been fully resolved. From our optogenetic experiments, we argue against a primary role of such activity in driving the reward-collection response. Further investigation is needed to understand the role of DLS representations of reward collection during operant behaviors.

Another point worth noting is that the engagement of DLS is time-sensitive. We consistently found larger effects on movement initiation and execution when the inhibition was closer to the onset of a specific movement transition, for example, right before paw lifting or immediately after paw lifting, but not toward the end of a reaching movement. Similarly, inhibition closer to the lever press onset impaired the pressing movement, whereas inhibition further from the lever press produced smaller effects. Following lever release, the turning movement was severely impaired, whereas inhibition coupled to the trigger stimulus also impaired the subsequent reward-retrieval process, but to a smaller extent (compare Fig. 7c–i to fig. S7d–j). This observation underscores the usefulness of temporally precise optogenetic perturbation for dissecting the moment-by-moment contributions of DLS activity to ongoing behavior.

### Comparisons with previous work

In head-fixed mice, striatal inactivation with archaerhodopsin-3 (ArchT) during joystick movement reduced movement velocity while leaving displacement unchanged (*22*). This observation is comparable to our experiment in which DLS photo-inhibition was applied during forepaw lift, causing animals to drop the paw mid-air and then re-initiate the reaching movement moments later (Fig. 5h). Notably, paw dropping was not reported in the joystick task, possibly because the joystick served as a support that prevented it. Consistent with the joystick result, when photo-inhibition was applied near the pressing phase (later in the reach), lever-press movements were impaired (Fig. 5l,m), suggesting impaired force generation during press execution when DLS activity was suppressed.

Studies using lever-press tasks have made observations supporting forelimb initiation deficits with DLS inhibition. In a study where striatal inactivation by ArchT was coupled to lever approach, the duration from approach to lever press increased, suggesting impaired initiation of lever pressing (*27*). Similarly, halorhodopsin-mediated DLS inhibition increased completion time on a fixed-ratio (FR-3) task, in which rats were trained to make multiple presses to earn a reward. This was likely because of initiation difficulty, as inter-press intervals increased (*63*). In our study, we observed an increase in approach-to-press duration and quantitatively analyzed the forepaw movement, identifying that the increased duration resulted from a delay in lift initiation rather than a general slowing of forelimb movement during lever reaching or difficulty pressing the lever (Fig. 4j). Moreover, we coupled the inhibition to ongoing reaching using online pose tracking, further testing whether DLS is also required for completing initiated movements (Fig. 5), an approach that, to our knowledge, had not been previously implemented.

Previous studies have reported reduced whole-body movement vigor following striatal inhibition or lesions. For example, in a decision-making task in a T-maze, DLS inhibition by halorhodopsin increased trial duration in rats (*64*). Similarly, rats with lesions of the dorsal striatum moved more slowly on a treadmill while learning to time their actions (*18*). In another study, inhibition of direct-pathway SPNs in mice performing an interval-categorization task increased “choice time” (*65*). “Choice time” was defined as the movement duration from a central port to a side port that delivered a reward when selected correctly. Thus, an increase in choice time suggests a reduction in whole-body movement vigor, consistent with our finding during reward retrieval (Fig. 7).

Our work extends these studies in several important ways. First, we used a robust method to inactivate DLS neurons, achieving a median of 90% suppression of spiking in putative DLS SPNs. Our approach did not require the opsin to be expressed in DLS SPNs, since inhibitory interneuron activation can inactivate nearby projection neurons broadly. Second, we performed side-by-side comparisons of the effect of DLS inhibition across distinct movement types, allowing us to establish the causal influence of DLS spikes on forelimb and whole-body movements, as well as the relative lack of effect on the nose poke movement. Third, we quantitatively tracked the effect of DLS inhibition on movement trajectories, a feature that has been limited or absent in previous studies. Fourth, by combining DLS photo-inhibition with SNr recordings, we directly linked striatal perturbations to changes in BG output, providing a circuit-level account of how continuous DLS activity shapes ongoing behavior. Finally, we compared the effects of DLS and DMS inhibition and found a clear dissociation in their behavioral consequences, with DLS inhibition strongly affecting forelimb initiation and execution, whereas DMS inhibition produced only mild slowing of locomotion.

### Limitations

Rats are well suited for reaction-time experiments: their forelimb and whole-body movements are quantifiable in freely moving paradigms, their lever-press behaviors become stereotyped once stabilized, and their size accommodates large implants with minimal impact on behavioral performance. However, our optogenetic approach, which drives inhibitory interneurons to suppress the firing rates of projection neurons (*41*, *66*), does not isolate cell-type- or pathway-specific effects, as both the direct (striatonigral) and indirect (striatopallidal) pathways are suppressed. Future work could leverage transgenic rats (*67*) or molecular tools (*68*) to perturb these pathways selectively. Alternatively, downstream circuits could be targeted directly, including the external globus pallidus (GPe) (*69*) or DLS outputs could be silenced using axonal-silencing approaches (*60*). The current approach and its variants (e.g., activating striatal PV-positive interneurons) produce robust suppression of SPN spiking, functionally akin to a precisely timed lesion, and can motivate more fine-scale perturbations. Moreover, pathway-specific manipulations are often complicated by lateral interactions: increasing or decreasing the activity of one cell type inevitably alters the activity of the other (*39*, *69*).

During the reward retrieval phase, rats executed a turning motion toward the reward port. DLS photo-inhibition markedly slowed this movement and altered the trajectory (Fig. 7d). One possibility is that turning, relative to straight-line locomotion, relies more heavily on the DLS, which could account for much of the observed slowing in whole-body movement. Turning is indeed strongly represented in the DLS (e.g., unit 7 in Fig. 2; unit 7 in fig. S3; see also (*11*, *28*)). However, our behavioral box was not large enough to fully assess straight-line locomotion; rats typically reached the reward port shortly after turning and began to decelerate. Future experiments in a maze (e.g., a T-maze (*9*, *64*, *70*)) that combines turning with straight-line locomotion in a long corridor could directly address this question.

### Motor control and submovements

Our current work has focused on establishing the causal effect of striatal spikes on movement generation using a transient-inactivation strategy. Previously, we performed DLS lesions in the same behavior (*19*). Together, these two manipulations provide complementary insight into striatal contributions to movement control. Transiently suppressing DLS spikes via optogenetic inhibition impairs multiple movement types: forelimb reaching is delayed or interrupted, and whole-body locomotion is slowed. Consistently, lesions also altered forelimb reaching behavior and persistently slowed whole-body locomotion (*19*). These results jointly support a role for the DLS in improving the quality of movements by facilitating their initiation and continuous progression.

Seemingly single, continuous movements, for example, a forelimb reach or whole-body turn, can be decomposed into multiple distinct phases or submovements (*29–31*, *71*). From this perspective, DLS spikes may be involved in chaining these movement segments into smooth trajectories with high efficiency and velocity. Inhibiting the DLS impairs these transitions, producing movement interruption (e.g., paw dropping) for forelimb movements and slower, more difficult turning for whole-body movements. The small impact on nose-poke movements, despite substantial time-locked DLS activity, suggests that not all operant movements rely equally on DLS and that, in some cases, DLS spiking may instead reflect reward-predictive signals that are only loosely coupled to the immediate control of movement segments.

## Materials and Methods

### Animals

All animal procedures were conducted in accordance with the animal care standards set forth by the US National Institutes of Health and were approved by the Institutional Animal Care and Use Committee of Peking University. Male adult Brown-Norway rats were obtained from Vita River Laboratories (Beĳing, China). Rats were older than three months and weighed 250–400 g prior to water restriction and behavioral training. Rats were housed individually on a 12-hour reverse light-dark cycle. During water restriction, water was earned during behavioral sessions; if intake was less than 12 mL, supplemental water was provided at least 3 hours after the session. Body weights were maintained at approximately 80% of their free-feeding level. Training was conducted six days per week, with unrestricted water access on the seventh day.

### Apparatus and Behavioral training

Behavioral training protocols were detailed in a prior study (*19*). Briefly, custom operant chambers integrated lever-based response modules (Med Associates, Fairfax, VT, USA) with Bpod state machines (Sanworks LLC, Rochester, NY, USA) to precisely control reward delivery and event timing. Within each chamber, a retractable response lever and reward port were positioned on opposite walls, separated by 32 cm (optogenetics experiments) or 35 cm (Neuropixels recordings). Behavioral programs were written in MedState Notation (MSN) by Marcelo Caetano and Mark Laubach. SRT behavioral performance was shaped progressively across six stages: (1) autoshaping; (2) lever press; (3) lever release; (4) lever hold (FP increased from 0 to 1.5 s) with a wide release response window (2 s); (5) hold with rapid release incentives (response window reduced from 2 to 0.6 s); and (6) full SRT schedule (0.75, 1.5 s FP, 0.6 s response window). The lever remained extended throughout the sessions (except for autoshaping), with trials self-initiated by lever presses. Correct responses triggered illumination of an LED within the reward port, cueing the delivery of a 60-μL, 10% (w/v) sucrose solution upon nose poke entry. Error responses (premature or late release) were penalized with a timeout (4–10 s), with the lever light and house light turned off (lever not retracted). The imperative trigger stimulus was a 250-ms tone (2 kHz, 70 dB SPL) delivered via a speaker mounted above the lever. An infrared LED, synchronized to tone onset, provided a signal for aligning video timestamps with behavioral events. A side-view camera (MER-160-227U3M, Daheng Imaging, Beĳing, China; 100 fps) was positioned externally to record forelimb kinematics during lever pressing. An overhead camera (MER-160-227U3M; 100 fps) captured top-view images of locomotor behavior.

In closed-loop optogenetic experiments, a transparent acrylic plastic block was placed directly in front of the lever to constrain the movement angle and posture of the animals, thereby facilitating the detection of lever-approach and paw-lifting behaviors. A 40-cm rectangular gap allowed access for lever pressing, reducing presses from extreme side angles.

### Stereotaxic surgery

Prior to surgery, rats were given unrestricted access to water for at least two days. Animals were anesthetized with isoflurane (induction: 4%; maintenance: 1–2% in O₂) and mounted in a stereotaxic frame (MTM-3; World Precision Instruments). Rectal temperature was maintained at 37.5 °C using a feedback-controlled heating pad. Eyes were lubricated with erythromycin eye ointment (Qilu Pharmaceutical, Jinan, China). Fur was removed using an electric clipper, and the scalp was sterilized with iodine tincture followed by 75% ethanol (3×). The scalp was incised along the midline, and the pericranium was reflected using a blunt scalpel. The skull surface was further cleaned with 3% hydrogen peroxide solution. Craniotomies were drilled at the target coordinates, and the dura was carefully incised using a 30-gauge needle prior to injection or implantation.

### Virus injection

Virus injections and optical fiber implantation (described below) were performed on separate days. Injections were performed using glass pipettes (Drummond Wiretrol II, Broomall, PA, USA) mounted on a custom-made injector system (Stereotaxic Injector System, Janelia) or a motorized microsyringe pump (UMP3; World Precision Instruments) coupled to a 10-μL precision syringe (701 RN; Hamilton, Reno, NV, USA) fitted with a removable 33-gauge, 12-mm needle (Hamilton). AAV2/9-mDlx-hChR2(H134R)-EGFP-WPRE-pA, or a control virus, AAV2/9-mDlx-EGFP-WPRE-pA (titer: 3×10^12^vg/mL; Shanghai Taitool Bioscience, Shanghai, China) was injected bilaterally into the dorsolateral striatum (relative to bregma; first site: AP = +1.3 mm, ML = ±3.6 mm; second site: AP = +0.3 mm, ML = ±4.0 mm) or dorsomedial striatum (first site: AP = +1.5 mm, ML = ±2.5 mm; second site: AP = +0.5 mm, ML = ±2.7 mm). Each site included injections at two depths (DV = −4.6 mm, −4.0 mm, relative to the brain surface), and 250–300 nL of virus was injected at each depth at a rate of 50 nL/min. After each injection, the pipette was left in place for 10 min before slow retraction. Craniotomies were sealed with Kwik-Cast silicone sealant (World Precision Instruments) after all injections at a site were completed. The scalp was sutured and rats were allowed to recover in their home cages.

Flunixin meglumine (2.5 mg/kg, subcutaneous or s.c.; QianShou Bio, Wuhan, China) was administered immediately post-surgery and daily for 3–5 days for analgesic effects, alongside ceftriaxone sodium (30 mg/kg, s.c.; Yuekang Duozhi, Beĳing, China) to prevent infection. Body weights were monitored daily until full recovery.

### Optical fiber implantation

After a minimum of 2 weeks of recovery, the scalp incision was reopened, and the skull surface was cleaned as described for virus injections. Four stainless-steel anchor screws (1.2 mm diameter; Yuyan Bio, Shanghai, China) were inserted into the skull for stability. Fiber-optic cannulae (200 μm diameter, NA = 0.37; InPer Technologies, Hangzhou, China) were bilaterally implanted in the DLS (AP +0.8 mm, ML ±4.0 mm, DV −3.9 mm relative to bregma) or in the DMS (AP +1.0 mm, ML ±2.3 mm, DV −3.9 mm). The craniotomies were sealed with Kwik-Cast, and the ferrules were secured to the skull using Super-Bond C&B dental adhesive (Sun Medical Co., Ltd., Shiga, Japan). A protective layer of dental cement (Jet Denture Repair; Lang Dental, Wheeling, IL, USA) was applied over the exposed skull and screws. Rats were allowed to recover under a heated lamp (35–37 °C) until fully ambulatory (15–30 min).

### Electrode implantation

Several types of electrode implants were used for DLS recordings across rats. Custom-made 16-channel fixed microwire arrays (MWAs, nickel-iridium, BioSignal, Nanjing, China) were implanted in four rats. For combined optogenetic and electrophysiological recordings in the striatum, 16-channel fixed MWAs (Hong Kong Plexon, China) integrated with a 200-μm optical fiber were implanted in three rats (fig. S2b). In two rats, 32-channel silicon probes (Diagnostic Biochips) were mounted on a movable drive (Micro Linear Drive, Janelia) and implanted. The coordinates of these electrode implants were reported previously (Figure 2 in Zheng et al. (*19*)). In two additional rats, an immobile Neuropixels probe 2.0 (IMEC, Leuven, Belgium) was implanted using a published protocol (*72*).

Surgical procedures followed standard stereotaxic methods, as described above. Two ground screws, each with a silver wire soldered to it, were implanted bilaterally in the skull above the cerebellum, ensuring contact with brain tissue for reference and grounding. Target electrodes were implanted contralateral to the preferred paw at the following coordinates (relative to bregma): AP = +0.8 mm, ML = 4.0 mm. Fixed arrays were inserted to a final depth of −4.0 to −4.5 mm (relative to the brain surface), with adjustments made during implantation when large-amplitude spikes (> 100 μV) were observed on the recording system. The silicon probe was lowered to an initial depth of 2.6 mm (relative to the brain surface) and subsequently advanced in 50 μm increments after each recording session. The Neuropixels probes, mounted on a reusable drive (3Dneuro) (*72*), were inserted to a depth of 5.5 mm (relative to the brain surface) at a 6° angle in both the anterior–posterior and medial–lateral planes and remained immobile thereafter. The craniotomy and the electrode were sealed with petroleum jelly (Vaseline) and bone wax. Neural recordings were performed after one week of post-implantation recovery.

For SNr recordings, movable tetrode arrays (Kedou Biotech, Hangzhou, China; stainless steel, 12.5-μm-diameter polyimide-coated wires) were used in an additional cohort of rats. These consisted of 16-channel (4 tetrodes) or 32-channel (8 tetrodes) configurations mounted on a microdrive. Target coordinates (relative to bregma) were: AP = −5.2 mm, ML = ±2.4 mm. Electrodes were initially lowered to a depth of −7.3 mm (relative to the brain surface) and then advanced in 50 μm increments after each recording session to optimize yield.

Multi-channel recordings from the MWA, tetrode, or silicon probes were conducted using a digital amplifier (Cereplex Direct, Blackrock Neurotech, Salt Lake City, UT, USA). Neuropixels recordings were conducted using a PXIe-based system (National Instruments, Austin, TX, USA) and acquired with the SpikeGLX software environment (Janelia Research Campus).

### Neural data analysis

#### Inclusion criteria

All chronic recordings were screened to exclude duplicate units recorded across multiple sessions. For Neuropixels data, 10–11 sessions per animal were included. An automated algorithm (Huang et al., *unpublished, preprint will be uploaded soon*), followed by manual curation, was used to identify and remove repeated units across sessions, ensuring that unique units were included in the population analysis. Moreover, units were excluded if > 2% of their spikes’ inter-spike intervals (ISIs) were < 3 ms. For all DLS units, the median ISI violation rate was 0.1% (IQR: 0–0.4%, *n*= 593 putative SPNs); for all SNr units, the median ISI violation rate was 0.1% (IQR: 0.1–0.5%, *n*= 300 SNr units).

#### Spike waveform analysis

Spike waveforms were inverted so that negative potentials were oriented upward (Fig. 1f). Spike width was quantified as the full width at half maximum (FWHM). To distinguish units with distinct waveform shapes, principal component analysis (PCA) was applied to the waveforms of all units within a given brain region (DLS or SNr), treating the voltage value at each time point as a feature. The average waveform of each unit was then projected into the resulting PCA space for low-dimensional visualization. A scatter plot of the projection onto the first principal component (PC1) versus FWHM was used to separate putative spiny projection neurons (SPNs) from non-SPNs. Units were clustered into two groups using the density-based method HDBSCAN (*73*). Unclustered data points were assigned to the nearest cluster based on proximity. Putative SPNs were identified as the cluster exhibiting broader spike widths (Figure 1g).

#### Spike rate modulation by behavioral events (GLM)

To investigate how behavioral events modulated neuronal spike rates during task performance, spike counts were quantified within pre- and post-event time windows relative to key behavioral events using a generalized linear model (GLM) (see fig. S2a). The events and their respective time windows were (in seconds): (1) approach/lift, [−2, −1] (pre) and [−1, 0] (post) relative to lever press; (2) press, [−0.5, 0] (pre) and [0, 0.5] (post) relative to lever press; (3) trigger/release, [−0.25, 0] (pre) and [0, 0.25] (post) relative to trigger stimulus; (4) poke, [−1, 0] (pre) and [0, 1] (post) relative to port nose poke. Events with fewer than 20 total spikes across trials were excluded to ensure reliable parameter estimation.

Spike count (spike_count_*ij*_) was treated as a random variable from a Poisson process: Poisson(μ_*ij*_). The goal was to determine whether μ was modulated by the event time point (post relative to pre) for each event. A GLM with a log-link was fitted to each event separately using the “fitglm” function (MATLAB, MathWorks). For an event, if either the pre- or post-window had zero counts in > 50% of trials, we added 1 to both windows to stabilize fitting. The model included a fixed effect for event time point (pre: 0; post: 1) and an offset for window duration:

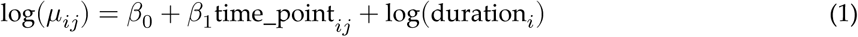

where the response variable spike_count_*ij*_ follows a Poisson(μ_*ij*_) distribution; *i* indexes the time window (pre: *i*= 1, post: *i*= 2) for trial *j*; β_0_ represents the fixed intercept; β_1_ is the fixed-effect coefficient for the event time point; time_point_*ij*_ is the indicator variable for pre or post time window (pre: 0, post: 1); and duration_*i*_ is the duration (in seconds) of the respective time window (e.g., 0.25 s for the trigger/release pre and post windows). The coefficient β_1_ represents the log-fold change in spike rate between post- and pre-event windows.

A contrast on β_1_ was evaluated using a two-sided Wald test (“coefTest” function) to test the null hypothesis: β_1_ = 0, yielding a *p*-value that indicates whether that unit’s spike rate changed between post- and pre-windows for that event. To control the false discovery rate (FDR) at 0.05, the Benjamini-Hochberg procedure (*74*) was applied across all units and events within a brain region (DLS or SNr) to derive a critical threshold; unit-event pairs with *p*-values below the threshold were deemed significantly modulated (cf. Fig. 1l).

#### SDF and warping

Spike times were convolved with a Gaussian kernel (σ = 50 ms) to obtain a smoothed spike density function (SDF). SDFs were aligned to event times to obtain a peri-event SDF, with 95% CIs derived from bootstrap resampling (1000 iterations). To average trials with different reaction times and reward retrieval durations, SDFs were linearly warped to target times (median reaction time or reward retrieval duration across all trials at the animal level; and across animals for population averages) (*75*). Let *t*=*t*_1_, *t*_2_, …, *t*_*n*_ be the time points of the SDF *f*(*t*) for the current trial, and let the target time points be *t*^′^ =*t*^′^, *t*^′^, …, *t*^′^. The scaling factor *s* was applied to warp the SDF, yielding *SDF*_warp_(*t*^′^), denoted as *f*_warp_(*t*^′^), where *t*^′^ =*t*^′^, …, *t*^′^, as follows:

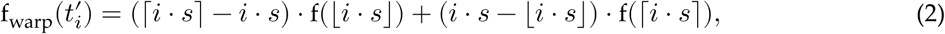

where 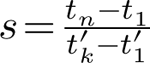 and ⌈⋅⌉ and ⌊⋅⌋ are the ceiling and floor functions, respectively.

Units were classified as positively modulated if the maximum of their mean-subtracted average SDF (computed from 750- and 1500-ms trials) exceeded the absolute value of the minimum; otherwise, they were classified as negatively modulated. For colormaps showing ranked activity of DLS or SNr neurons (Figures 1k and 9j), the warped SDF, *SDF*_warp_(*t*), was mean-subtracted and soft-normalized by division by (*r*+*c*), where *r* was the min-max range and *c* was a constant (*c* = 5)(*76*). To order units by response timing, data from correct trials in the 750 ms-FP condition were used to order units by the first time each unit’s mean-subtracted SDF crossed a threshold (75% of the maximum for positively modulated units or 75% of the minimum for negatively modulated units). The same order was then applied to the 1500 ms-FP condition.

#### PCA

Neural population activity was analyzed using PCA to identify low-dimensional manifolds that capture the dominant patterns of variance across units. Unlike prior work (*19*), to avoid mixing different behavioral regimes, PCA spaces were constructed separately for periods dominated by forelimb movements (lever press, hold, release; from 2.5 s before lever press to lever release) and whole-body movements (reward retrieval; from lever release to 1 s after nose poke). For each of the two behavioral epochs, soft-normalized SDFs from included units were stacked into a matrix of size time points × units, and PCA was then applied using MATLAB’s “pca” function, treating each unit (column) as a covariate, yielding principal components (PCs) ordered by explained variance. Population activity was then projected onto the top four PCs for visualization (Fig. 1l), which revealed that individual PCs aligned with distinct behavioral features, such as forelimb lift initiation or reward retrieval movement.

To identify subpopulations contributing significantly to behaviorally coupled PCs (Figure 1m), units were selected based on the magnitudes of their PCA loadings (weights *w*_*i*_; *i*= 1, 2, 3, … indexes the PCs) in the coefficient matrix. Thresholds were empirically pre-specified for different PCs. Loadings were taken from PCA on activity aligned to lever press for the lift/hold/trigger phases, and to nose poke for retrieval:

1. *Lift* units (selective for pre-press paw-lifting movement): DLS: (*w*_2_ > 0.05) SNr (positively modulated): (*w*_2_ > 0.05) SNr (negatively modulated): (*w*_2_ < −0.05)
2. *Hold* units (selective for sustained press): DLS: (*w*_1_ > 0.05) SNr (positively modulated): (*w*_1_ > 0.05) SNr (negatively modulated): (*w*_1_ < −0.05)
3. *Trigger/Release* units (selective for trigger stimulus and lever release): DLS: (*w*_2_ < 0.025 and *w*_3_ > 0.05) SNr (positively modulated): (*w*_3_ < −0.05) SNr (negatively modulated): (*w*_3_ > 0.05)
4. *Retrieval* units (selective for post-release reward retrieval): DLS: (*w*_1_ <−0.075 or *w*_2_ > 0.075 or *w*_3_ > 0.075) SNr (positively modulated): (*w*_1_ <−0.075 or *w*_2_ > 0.075 or *w*_3_ <−0.075) SNr (negatively modulated): (*w*_1_ > 0.075 or *w*_2_ <−0.075 or *w*_3_ > 0.075)

This resulted in partially overlapping subsets of units per phase, each enriched for functional relevance to the associated behavioral events. The overlaps are illustrated in the Venn diagrams (fig. S2d,g). Substantial overlap indicates that a unit showed multi-event modulation.

#### Mapping firing rate to movement trajectories

Following alignment of video frames to behavioral timestamps, video clips were extracted around each lever press: −5 to +5 s for top-view videos, and −2 to +2 s for side-view videos. Body-part pose models were trained on selected frames and videos were tracked using DeepLabCut (*37*). Tracked body parts were then manually inspected and curated via a custom MATLAB GUI to ensure high-quality trajectories (e.g., excluding occluded segments). Head position was defined as the midpoint between the two ears. For each video clip, the *x*−*y* coordinates of the tracked parts were stored in a table with each row representing a frame.

Discrete spike times were convolved with a Gaussian kernel (σ = 50 ms) to obtain continuous SDF (in Hz). The tracking arena was discretized into spatial bins: 1 × 1 mm for top-view trajectories and 0.33 × 0.33 mm for side-view trajectories. For each trajectory segment, head positions were linearly interpolated to the SDF’s timestamps. For each spatial bin, occupancy time was calculated as the total time during which the interpolated head position fell within the bin boundaries. The firing rate was estimated from the integral of the SDF over time (*C*_*b*_) and the occupancy duration (*T*_*b*_) in bin *b*, aggregated over all trials (Fig. 2g; fig. S3g). A 2-D Gaussian kernel (σ = 1 pixel) was applied for smoothing:

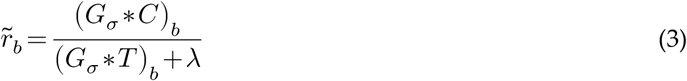

where (*G*_σ_ ∗*C*) denotes the 2-D convolution of *C* with the Gaussian kernel, evaluated at bin *b*; λ= 100 ms is a ridge constant that stabilizes estimates in unvisited or sparsely visited bins; and *r*∽_*b*_ is the estimated firing rate when the rat traversed bin *b*.

### Optogenetic stimulation during lever approach

#### Hardware and experiment pipeline

An infrared sensor (FQD-31NO, Luo Shida Sensor, Dongguan, China) was positioned above the lever to detect the approach of the rats. Once detected, in a small proportion of trials (<10%), laser stimulation (473 nm) was triggered. The stimulation consisted of a 2.2-s train at 40 Hz with 5-ms pulses (DPSS laser, MBL-FN-473, Changchun New Industries Optoelectronics Technology Co., Ltd., Changchun, China). Laser light was delivered to the brain through an optical fiber (250 μm; NA = 0.37). Laser power was calibrated at the tip each day before the experiment and was set to 20 mW peak, yielding an average power of 4 mW during the pulse train. To minimize visual confounds, a masking blue LED flash (20-Hz, 5-ms pulses for 4–6 s), triggered upon approach, was presented in both NoStim and PhotoInh trials.

Forelimb movement was recorded using high-speed video at 100 fps (camera: MER-160-227U3M, Daheng Imaging, China; acquisition: StreamPix 7, NorPix, Montreal, Canada). Post-acquisition alignment was performed using an infrared signal locked to the trigger stimulus. Forelimb movement was tracked using DeepLabCut (*37*).

A polyvinylidene fluoride (PVDF) vibration sensor (LDT0-028k, TE Connectivity) was affixed to the end of the lever. The electrical signal was amplified with an amplifier module (PVA103, Shenzhen Vkinging Electronics Co., Ltd, China), digitized with the multifunction data acquisition device PCIe-6341 (National Instruments), and stored using WaveSurfer (Janelia). Using simultaneously recorded video-derived lever position, a linear model converted the sensor’s voltage signal to lever position (*R*^2^ > 0.98 on testing data; fig. S10).

#### Approach-to-press duration analysis (***t*_a2p_**; Fig. 4h)

The approach-to-press duration (*t*_a2p_) was computed per trial as the elapsed time between the approach detection and the lever press time recorded by Bpod. Outliers were excluded if |*t*_a2p_ −median(*t*_a2p_)| > 5×MAD(*t*_a2p_). The effect of DLS photo-inhibition on the total duration from approach detection to lever press (*t*_a2p_) was evaluated using a linear mixed-effects (LME) model (*t*_a2p_ ∼ StimType + (1 + StimType | anm)), which included stimulation type (StimType, 0 for NoStim and 1 for PhotoInh) as a fixed effect, with random intercepts and slopes for StimType varying by animal (anm) to account for individual variability. The model is:

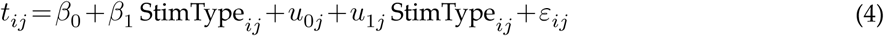

where *t*_*ij*_ denotes *t*_a2p_ for trial *i* in animal *j*, β_0_ is the fixed intercept (baseline for NoStim), β_1_ is the fixed effect of StimType, *u*_0*j*_ and *u*_1*j*_ are random intercepts and slopes for animal *j*, and ε_*ij*_ is the residual error. Random effects are assumed to follow a multivariate normal distribution: (*u*_0*j*_, *u*_1*j*_) ∼*N*(0, Σ), where Σ is the variance-covariance matrix, and residuals are normally distributed: ε_*ij*_ ∼*N*(0, σ^2^). The same assumptions were used for the other LME models.

#### Velocity analysis (Fig. 4i)

Forepaw velocity during the lifting phase was analyzed in a subset of rats (*n*= 16) with unobstructed forepaw tracking. Velocity was defined as the average speed (cm/s) over the trajectory from 3.5 cm below the lift-peak position to the lift-peak position (total path length divided by time). An LME model (Velocity ∼ StimType + (1 + StimType | anm)) was fitted with StimType as a fixed effect and random intercepts and slopes for StimType by animal:

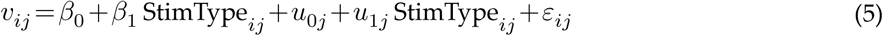

where *v*_*ij*_ is the observed velocity for trial *i* in animal *j*.

#### Segmented duration analysis (*t*_a2pk_ and *t*_pk2p_; **Fig. 4j**)

To dissect *t*_a2p_, each trial’s *t*_a2p_ was split into two segments: from approach to lift peak (*t*_a2pk_) and from lift peak to lever press (*t*_pk2p_). Segment type was treated as a factor (DurationType: a2pk or pk2p). An LME model (duration ∼ StimType × DurationType + (1 + StimType | anm) + (1 | anm:trial)) was fitted to assess the interaction between StimType and DurationType, with a random intercept and StimType slope by animal, plus a trial-level random intercept (anm:trial) to account for two repeated measures within a trial:

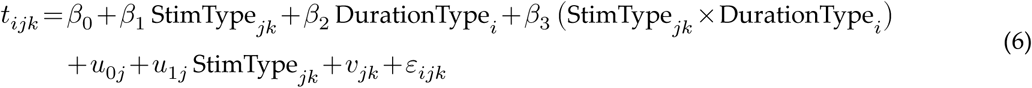

where *t*_*ijk*_ is the duration for DurationType *i* in animal *j* and trial *k*; *u*_0*j*_, *u*_1*j*_ are the animal-level random intercept and random slope for StimType; and *v*_*jk*_ is the random intercept per trial *k* within animal *j*.

#### Effect of different stimulation sides on ***t*_a2p_** (fig. S4e)

To examine the effects of photo-inhibition applied to different stimulation sides (bilateral, contralateral, or ipsilateral; bilateral as reference) on *t*_a2p_, an LME model was fitted with StimType and StimSide as fixed effects, along with their interaction: *t*_a2p_ ∼ StimType × StimSide + (1 + StimType | anm). Random intercepts and slopes for StimType varied by animal (anm). The model equation is:

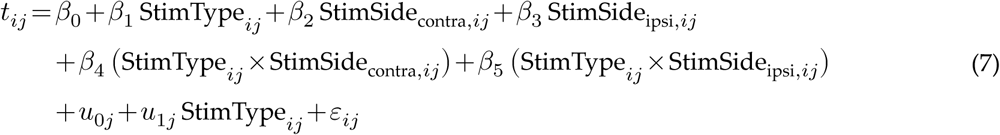

where *t*_*ij*_ is *t*_a2p_ for trial *i* in animal *j*.

### Optogenetic stimulation during forepaw lifting

A second side camera (MER-160-227U3M, Daheng Imaging, China) positioned next to the main side camera captured the paw-lifting process within a 20 × 15 cm field of view at 50 fps. Frames were streamed to Bonsai (*44*), where a smaller 7 × 4 cm region was analyzed. Bonsai integrated DeepLabCut-Live (*43*) to estimate the coordinates and detection likelihood of the selected forepaw (left or right). When the paw position exceeded a predefined threshold and the detection likelihood was > 75% for three consecutive frames, Bonsai transmitted a trigger signal to the Bpod state machine as a SoftCode event. Bpod then determined whether to initiate laser stimulation based on a predefined probability (< 10%). The threshold was adjustable to allow stimulation at different phases of the reaching movement (Fig. 5j–o).

#### Drop ratio analysis (Fig. 5g)

A drop event was counted when the vertical paw position fell below a predefined threshold (50–70% of the height from baseline to the global peak position) after the lift’s local peak.

The drop ratio (*p*_drop_) was computed for each condition (StimType per animal). Paired differences between NoStim and PhotoInh conditions were assessed using a two-sided Wilcoxon signed-rank test on the *p*_drop_.

#### Lift-to-press duration analysis (***t*_l2p_**; Fig. 5i)

The effect of DLS photo-inhibition on the duration from lift detection to lever press (*t*_l2p_) was evaluated using a LME model: *t*_l2p_ ∼ StimType + (1 + StimType | anm). The model included stimulation type (StimType, 0 for NoStim and 1 for PhotoInh) as a fixed effect, with random intercepts and slopes for StimType varying by anm:

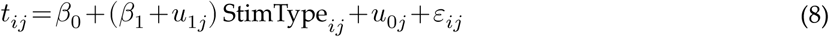

where *t*_*ij*_ is the *t*_l2p_ for trial *i* in animal *j*.

#### Segmented duration analysis (*t*_l2pk_ and *t*_pk2p_; **Fig. 5i**)

To dissect *t*_l2p_, each trial’s *t*_l2p_ was split into two segments: from lift to lift-peak (*t*_l2pk_) and from lift-peak to lever press (*t*_pk2p_). Segment type was treated as a factor (DurationType factor: l2pk or pk2p). An LME model (duration ∼ StimType × DurationType + (1 + StimType | anm) + (1 | anm:trial)) was fitted to assess the interaction between StimType and DurationType:

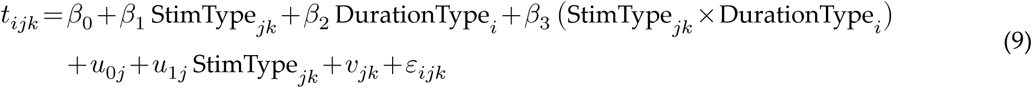

where *t*_*ijk*_ is the duration for DurationType *i* in animal *j* and trial *k*.

#### Peak-to-press duration at different stages (Fig. 5o)

The effect of photo-inhibition on *t*_pk2p_ was examined at different inhibition stages along the reach trajectory. The inhibition stage was treated as a factor Stage (0 for inhibition early in a reaching movement or 1 for inhibition late in a reaching movement). An LME model (duration ∼ StimType × Stage + (1 + StimType | anm)) was fitted to assess the interaction between StimType and Stage:

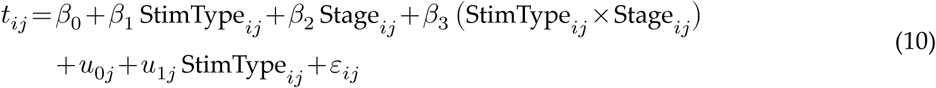

where *t*_*ij*_ is the *t*_pk2p_ for trial *i* in animal *j*.

#### Effect of different stimulation sides on ***t*_l2p_** (fig. S4j)

To examine the effects of photo-inhibition applied to different stimulation sides (bilateral, contralateral, ipsilateral) on *t*_l2p_, an LME model was fitted: *t*_l2p_ ∼ StimType × StimSide + (1 + StimType | anm). The model equation is:

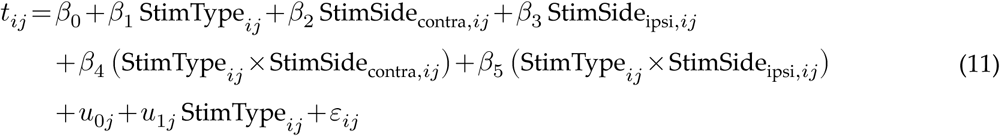

where *t*_*ij*_ is *t*_l2p_ for trial *i* in animal *j*.

### Optogenetic stimulation coupled to trigger stimulus

For trigger-coupled photo-inhibition (Fig. 6), the trigger stimulus activated the Bpod’s scheduled waves. Laser stimulation was applied in < 25% of trials per session, randomly assigned to different events, such as approach, lever-press, or trigger, within the same session.

Reaction-time (RT) outliers exceeding 25 × MAD from the median (> 4.427 s) were excluded. RTs < 0.1 s were also excluded. Photo-inhibition trials (and corresponding NoStim trials) were classified as ‘Early’ (≤ 60 stimulation trials) or ‘Late’ (> 60) to define the fixed factor StimGroup. FP is a categorical factor with two levels: Short = 750 ms and Long = 1500 ms. Log-transformed RT (logRT) data were fitted with an LME model (logRT ∼ StimType × StimGroup × FP + (1 + StimType | anm)):

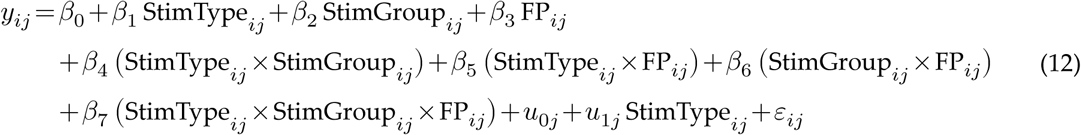

where *y*_*ij*_ is logRT for trial *i* in animal *j*.

### Reward retrieval duration and locomotor behavior

A top-view camera operating at 100 fps captured videos with StreamPix 7 to monitor locomotion. DeepLabCut was used to track body parts and determine the rat’s head location and direction. Movement paths were derived from the positions of the ears, with head direction defined as the vector orthogonal to the ear-to-ear line and pointing toward the nose. Instantaneous velocity of the head for each frame was calculated as the Euclidean displacement between the current head location and the head location one frame ahead, divided by the time interval (10 ms). The velocity time series was smoothed with a moving-average filter (centered window of 25 samples, or 250 ms). In release-coupled photo-inhibition, the mean velocity for each trial was computed as the average instantaneous velocity from lever release to 1 s afterward, capturing the period of active movement. For near-poke-coupled photo-inhibition triggered by an IR detector near the reward port, mean velocity was computed as the average from detection to 0.5 s afterward. Movement path length was calculated as the cumulative Euclidean distance from lever release (or IR detection) to the water-port nose poke. Photo-inhibition was applied bilaterally or unilaterally (ipsilateral or contralateral to the turning direction) in a randomized, interleaved order across sessions.

These release- or near-poke-coupled photo-inhibition experiments had a smaller sample size (*n*= 6) than the forelimb reaching experiments (*n*= 25 tested, *n*= 16 tracked), as the initial focus was on the forelimb movement. Whole-body locomotion tests were added later after observing prolonged reward retrieval duration under trigger-coupled photo-inhibition.

Three LME models with the same formula were applied to analyze movement path length, velocity, and retrieval duration. Measurements were obtained across multiple trials and sessions for each rat. Each LME model (Metric ∼ StimType × StimSide + (1 + StimType | anm)) included StimType and StimSide (bilateral, ipsilateral, contralateral to the turning direction; with bilateral as the reference level) as fixed effects, and animal as a random effect. The slope for StimType was allowed to vary among rats. The model for each measurement *y* is given by:

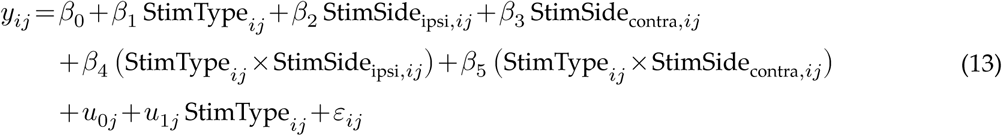

where *y*_*ij*_ is the metric (path length, mean velocity, or retrieval duration) for trial *i* in animal *j*; all other terms are defined as in the previous models. The exact model specification applies to both release- and near-poke-coupled photo-inhibition (Fig. 7g–i, n–p).

A similar model was fitted for reward retrieval duration under trigger-coupled photo-inhibition (fig. S7c). A simplified model excluding StimSide (bilateral only) was used for kinematics analysis during trigger-coupled photo-inhibition (fig. S7h–j).

### Statistics

LME or linear models were analyzed using R (“lme4” and “lmerTest” packages). Significance of the fixed effect (e.g., β_1_ in Equation 4) was assessed using a *t*-test with Satterthwaite approximation for degrees of freedom. Estimated marginal means for the PhotoInh - NoStim contrast (effect size) were computed using the “emmeans” package, with SEM derived from the model. *p*-values were adjusted using Holm’s method when multiple tests were performed (e.g., relative effect size among different StimSide conditions; fig. S6f–h). Effect sizes were reported for all statistical tests, typically as estimated marginal mean differences (PhotoInh − NoStim) with SEM. GLM-based analysis of spike count modulation was conducted using MATLAB, as detailed above.

## Acknowledgements

We thank Dr. Mark Laubach for sharing code for behavioral training and for many fruitful discussions. We thank Qiang Zheng for assistance with implementing real-time pose tracking.

## Author contributions

Conceptualization: JC, JY Methodology: JC, JY Investigation: JC, YZ, JY Visualization: JC, YZ, JY Supervision: JY Writing—original draft: JY Writing—review & editing: JC, JY

## Data and materials availability

All data and code supporting the results of this study will be made publicly available in an online repository upon publication. During peer review, data and analysis code are available on Dryad.

## Supplemental Materials

**Fig. S1.**
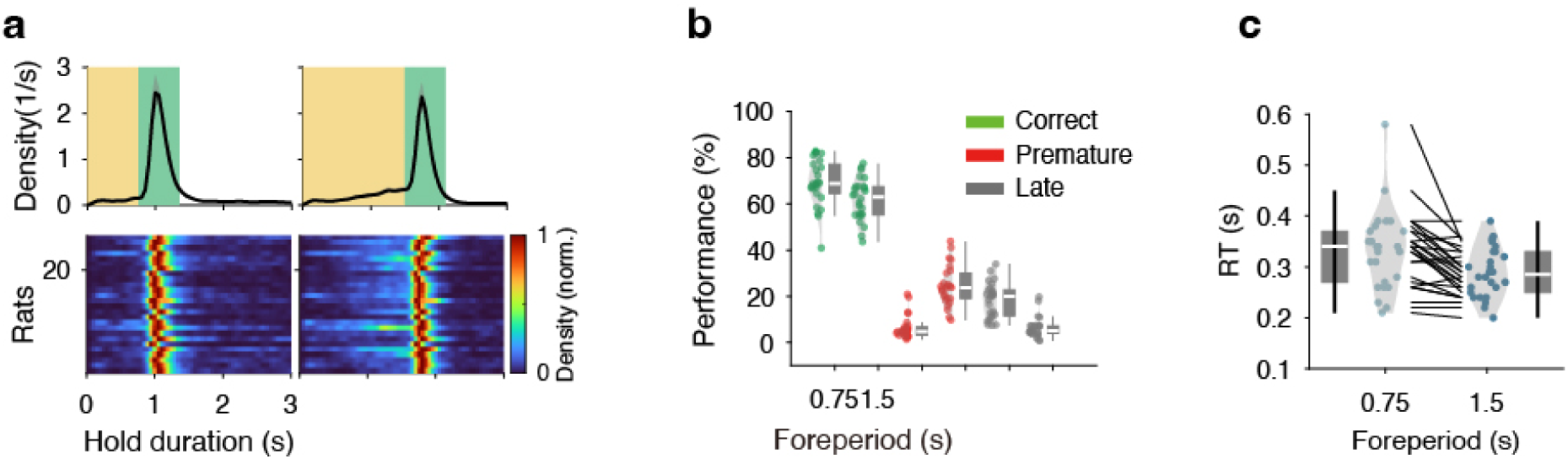
SRT behavior of rats. **a**, Probability density functions (PDFs) of 26 rats. Top, mean and SEM of PDFs for two FPs (left, 0.75 s; right, 1.5 s). Bottom, each row is the PDF of a single rat. Color denotes probability density (peak-normalized). **b**, Correct, premature, and late ratios. **c**, Reaction times of all rats for two FPs.

**Fig. S2.**
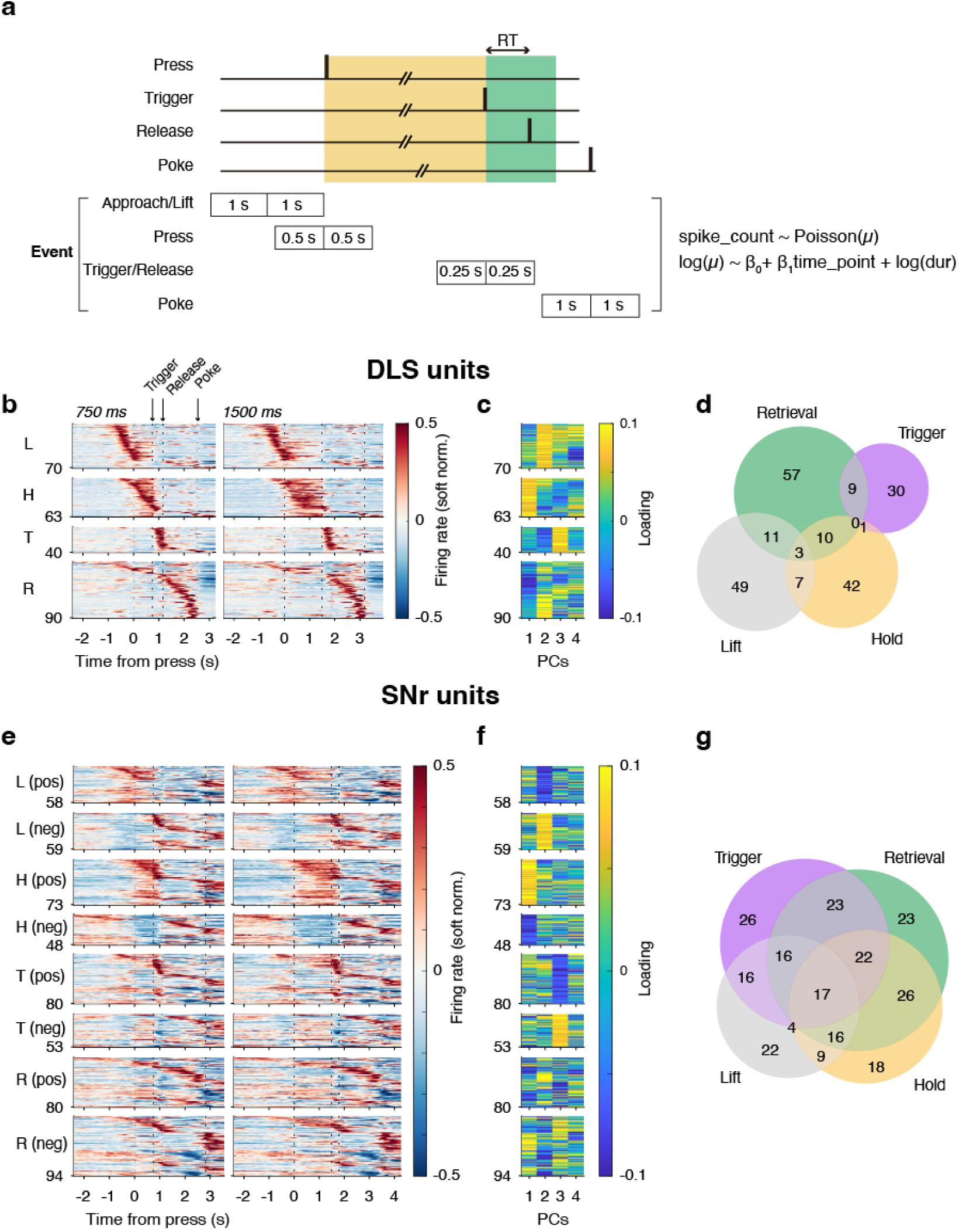
Behavioral event-modulation of neural activity in DLS and SNr. **a**, Diagram illustrating the construction of event time windows for analyzing spike rate modulation by a GLM. **b**, Left, Colormap showing the normalized SDFs of units selected by PCA loadings in Figure 1n. L: Lift; H: Hold; T: Trigger/Release; R: Retrieval. **c**, Projection loadings of these selected units. **d**, Venn diagram illustrating the overlap of selected units. **e–g**, Same as **b–d** for SNr units. Positively and negatively modulated units are shown separately for each functional category.

**Fig. S3.**
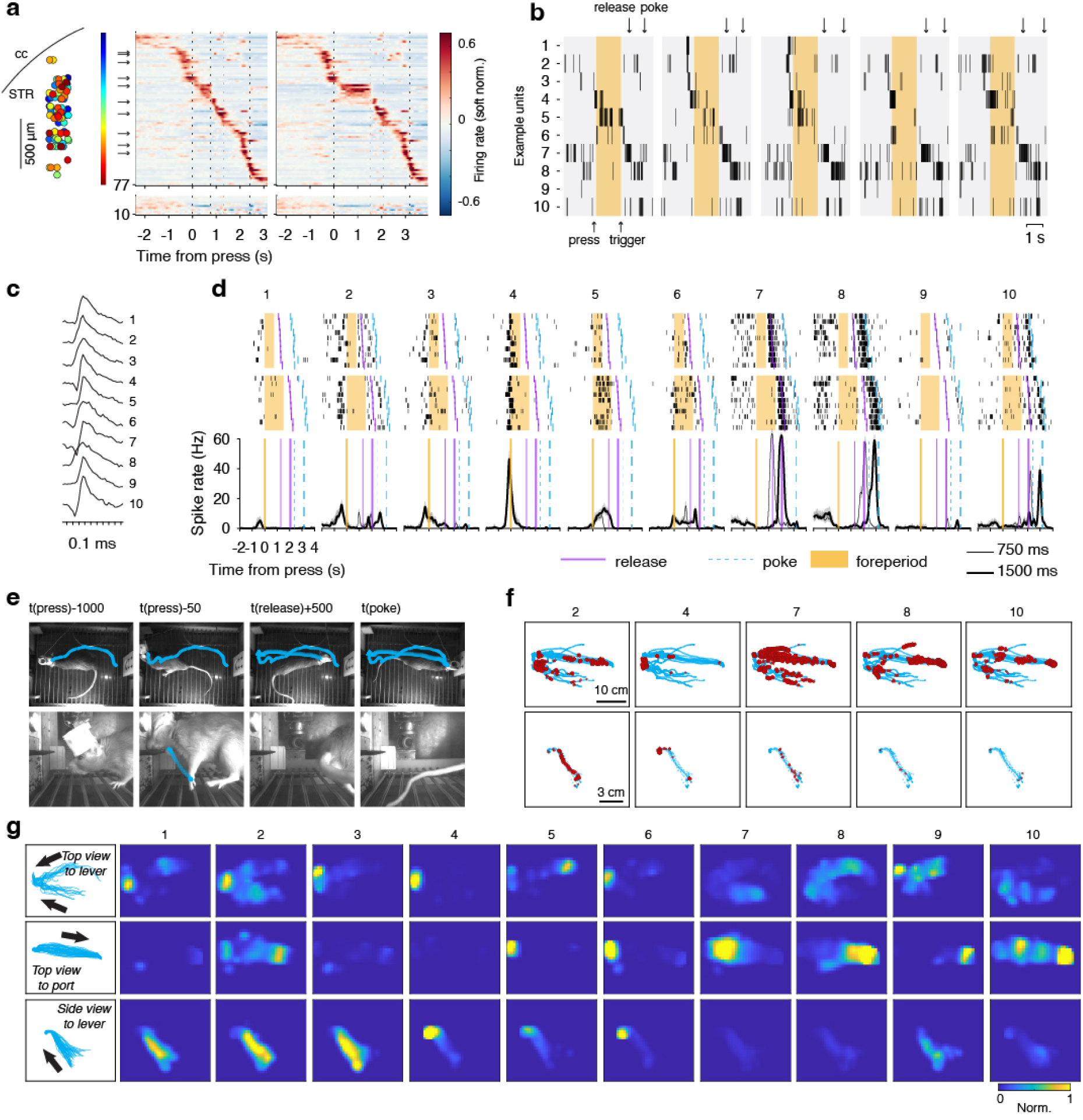
Striatal neural activity tracks rat’s movement in a different rat. As in Fig. 2 from chronic Neuropixels probe recordings in a different rat. All data are simultaneously recorded units from a single session.

**Fig. S4.**
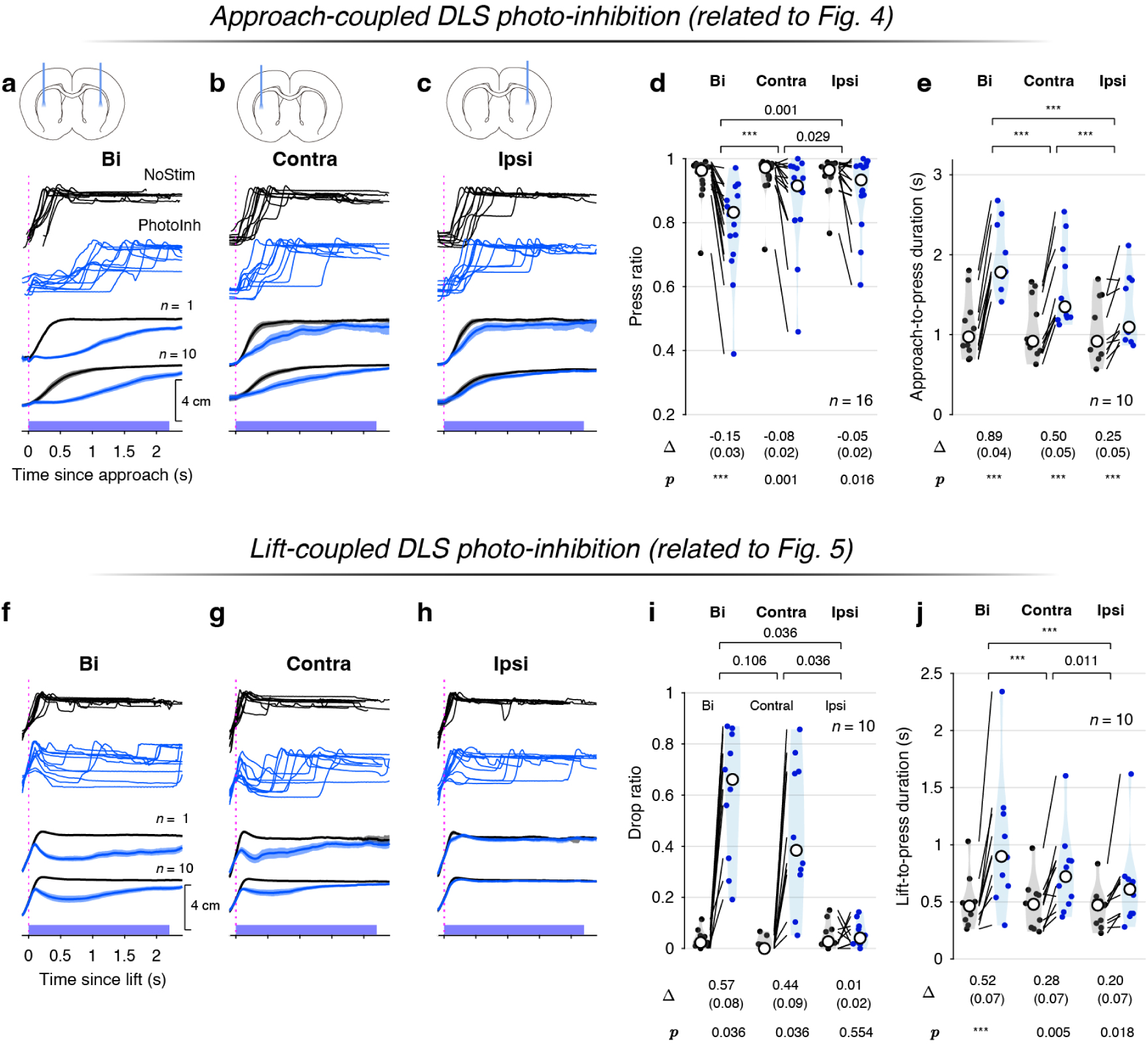
Unilateral versus bilateral DLS photo-inhibition during approach or lift. **a**, Same as Fig. 4d (approach-coupled stimulation) for a rat with bilateral (Bi) DLS photo-inhibition (*n*= 1) and the average response of all rats (*n*= 10). In these rats, both bilateral and unilateral (**b,c**) photo-inhibitions were performed and movement trajectories tracked. For each panel, the first three rows illustrate an example rat (final row: mean and 95% confidence intervals). The fourth row shows the mean and SEM of 10 rats. **b**, Same as **a** for trials in which DLS photo-inhibition was applied to the side contralateral (Contra) to the lifting paw. **c**, Same as **a** for trials in which DLS photo-inhibition was applied to the side ipsilateral (Ipsi) to the lifting paw. **d**, Violin plots display press ratio for no stimulation (NoStim, black) and DLS photo-inhibition (PhotoInh, blue) conditions across bilateral, contralateral, and ipsilateral stimulation sides, with lines connecting paired data from individual rats (*n*= 16 rats). Mean changes (PhotoInh −NoStim), SEM (in parentheses), and *p*-values from two-sided Wilcoxon signed-rank tests are listed below each stimulation side. To compare stimulation-induced changes across sides (e.g., Bi vs Contra), a Friedman test was conducted on the (PhotoInh−NoStim) differences, treating animals as blocks (Friedman χ^2^ = 16.625, d.f. = 2, *p* = 0.0002). Post-hoc pairwise Wilcoxon signed-rank tests were applied and Holm-adjusted *p*-values are shown above the violin plots between pairs of stimulation sides. ∗∗∗denotes *p* < 0.001. **e**, Violin plots display approach-to-press duration (*t*_a2p_) distributions (see Equation 7). **f–h**, Same as Fig. 5d (lift-coupled stimulation) for a rat with bilateral (Bi) DLS photo-inhibition (*n*= 1) and the average response of 10 rats (*n*= 10) for which both bilateral (**f**) and unilateral (**g,h**) photo-inhibitions were performed. **i**, Same as **d** for drop rate when DLS photo-inhibition was applied during lifting. To compare the effect across different stimulation sides, the Friedman test was conducted (Friedman χ^2^ = 16.2, d.f. = 2, *p* = 0.0003). **j**, Same as **e** for lift-to-press duration (*t*_l2p_ (Equation 11).

**Fig. S5.**
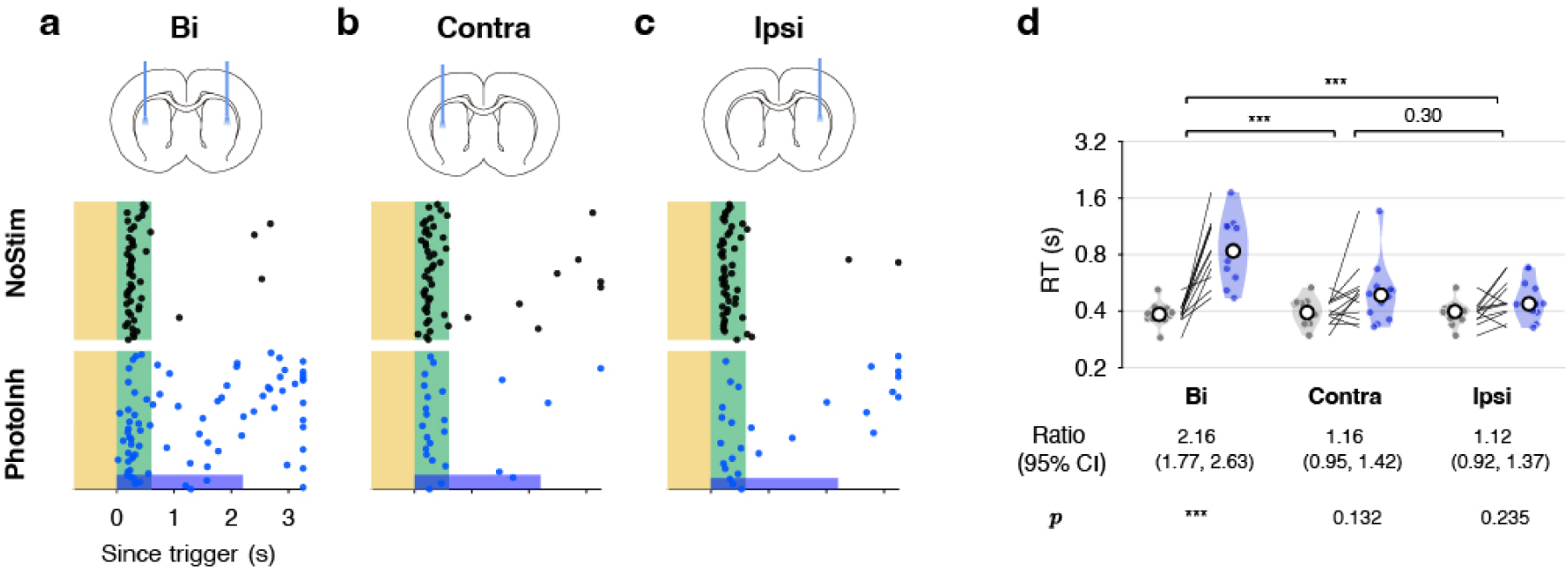
Unilateral versus bilateral DLS photo-inhibition coupled to trigger stimulus. **a–c**, Lever-release responses in an example rat during bilateral (Bi, **a**), contralateral (Contra, **b**), and ipsilateral (Ipsi, **c**) DLS photo-inhibition. Top row, NoStim; bottom row: PhotoInh. **d**, Geometric mean reaction time (RT) ratios (PhotoInh/NoStim) across rats under Bi, Contra, and Ipsi conditions. Data were analyzed using a LME model on log-transformed RTs (log(RT) ∼ StimType × StimSide + (1 + StimType ∣ anm)), where StimType (PhotoInh or NoStim) and StimSide (Bi, Contra, or Ipsi) are fixed effects. EMMs were computed for each combination, followed by post-hoc contrasts testing the log(RT) differences (PhotoInh−NoStim) within each StimSide and across StimSides (Bi - Contra, Bi - Ipsi, Contra - Ipsi), with Holm-adjusted *p*-values. Ratios and 95% confidence intervals are exponentiated to the original RT scale; ∗∗∗ denotes *p* < 0.001.

**Fig. S6.**
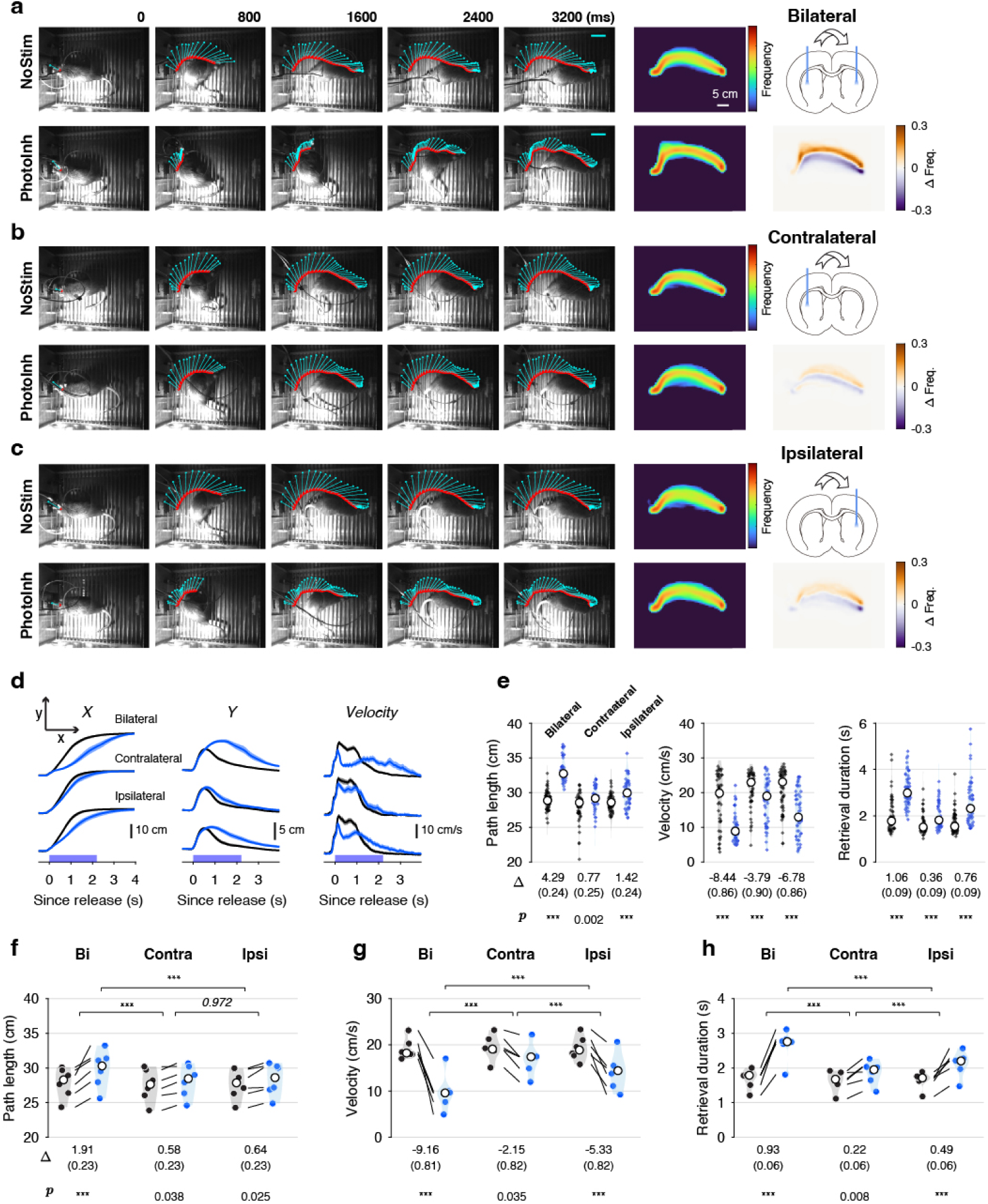
Unilateral versus bilateral DLS photo-inhibition during reward retrieval. **a**, Movement trajectories and frequency map differences for bilateral photo-inhibition. Arrows above coronal brain sections indicate the turning direction toward the reward port post-lever release; fiber locations show photo-inhibition sites. Color maps indicate pixel traversal frequency differences (PhotoInh −NoStim). **b**, As in **a** for contralateral photo-inhibition. **c**, As in **a** for ipsilateral photo-inhibition. **d**, Mean movement in X and Y dimensions and instantaneous velocity with 95% confidence intervals across stimulation sides (bilateral, ipsilateral, contralateral) for a representative rat. **e**, Movement path length, mean velocity, and retrieval duration for the representative rat; mean differences (SEM in parentheses) and Holm-adjusted *p*-values from a linear model shown below. **f–h**, As in **e** for *n*= 6 rats; Holm-adjusted *p*-values are from post-hoc pairwise contrasts based on EMMs from the LME model; ∗∗∗ denotes *p* < 0.001.

**Fig. S7.**
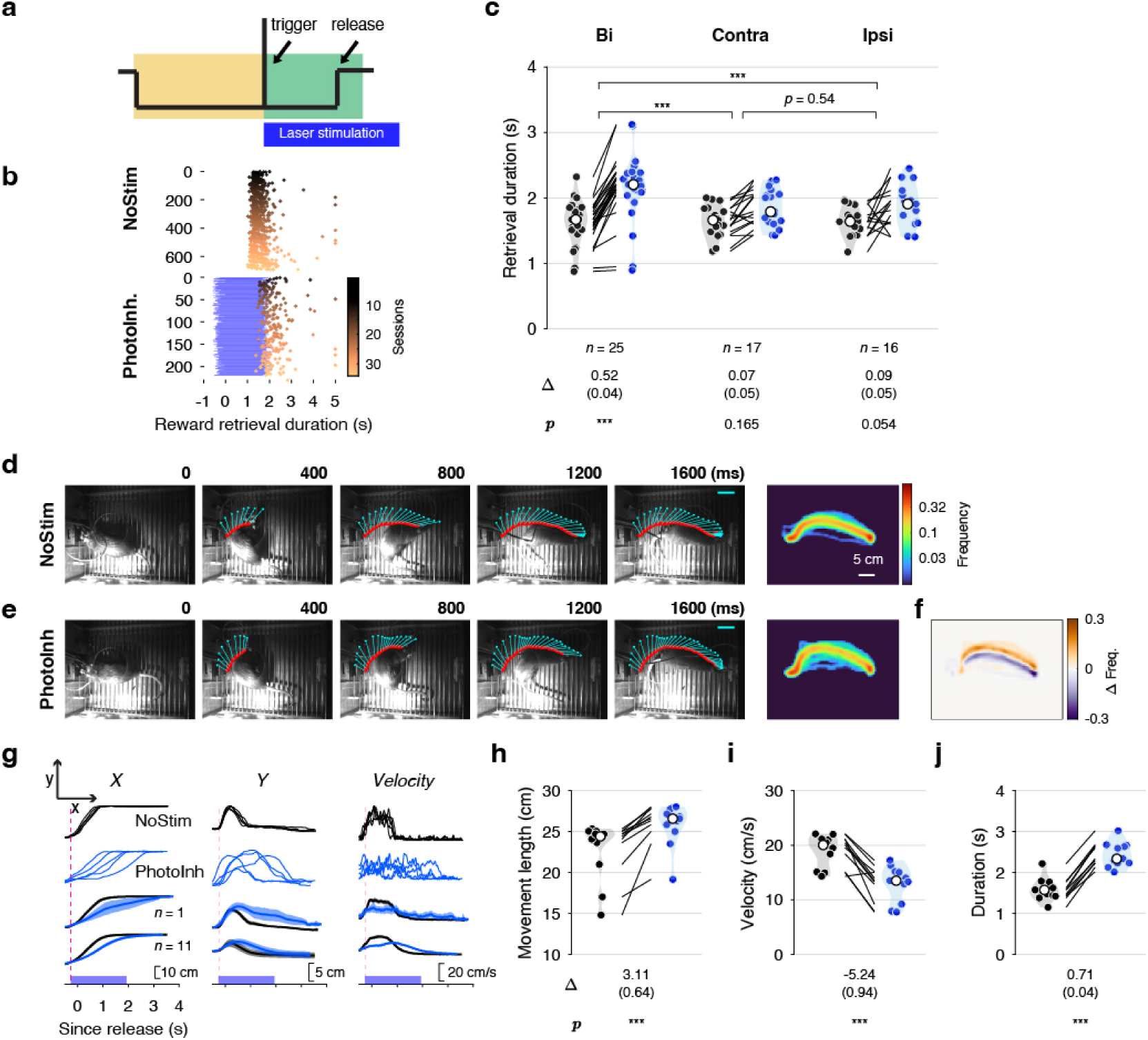
Trigger-coupled DLS photo-inhibition slows down reward retrieval and locomotion. **a.** Schematic of trigger-coupled DLS photo-inhibition. **b.** Reward retrieval duration with and without bilateral photo-inhibition (25% of trials) in a representative rat. Each dot represents duration to reach the reward port after lever release (time = 0); blue shading indicates photo-inhibition duration. **c.** Mean differences in retrieval duration (SEM in parentheses) for bilateral or unilateral photo-inhibition; Holm-adjusted *p*-values are from post-hoc pairwise contrasts based on EMMs from a LME model (duration ∼ StimType × StimSide + (1 + StimType ∣ anm)). **d–f.** As in Fig. 7c–e for photo-inhibition starting at trigger stimulus onset; time 0 denotes lever release. **g.** Movement in X and Y dimensions and instantaneous velocity; blue rectangles indicate photo-inhibition period aligned to trigger onset. **h–j.** Mean movement path length, mean velocity, and retrieval duration in *n*= 11 rats; Holm-adjusted *p*-values are from post-hoc pairwise contrasts based on EMMs from a LME model; ∗∗∗ indicates *p* < 0.001.

**Fig. S8.**
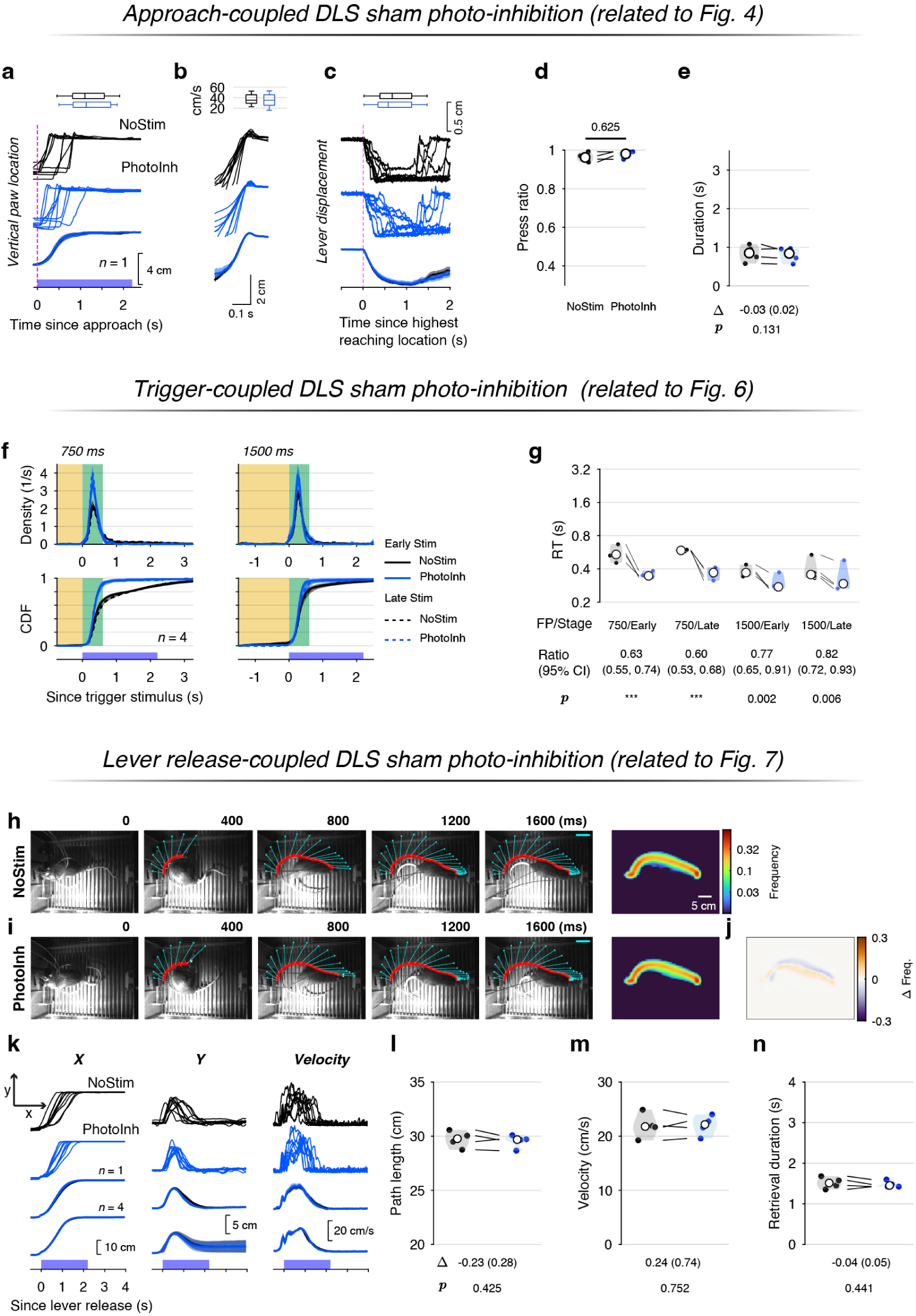
Control experiments. **a–e**, As in Fig. 4d–h but for control group (*n*=4 rats) expressing mDlx-GFP (instead of mDlx-ChR2-GFP) in the DLS. Experiments (DLS photo-stimulation was coupled to lever approach) and statistical analyses (with LME) matched the photo-inhibition group. **f–g**, As in Fig. 6c,d for control rats. DLS photo-stimulation was coupled to trigger stimulus. **h–n**, As in Fig. 7c–i for control rats. DLS photo-stimulation was coupled to lever release.

**Fig. S9.**
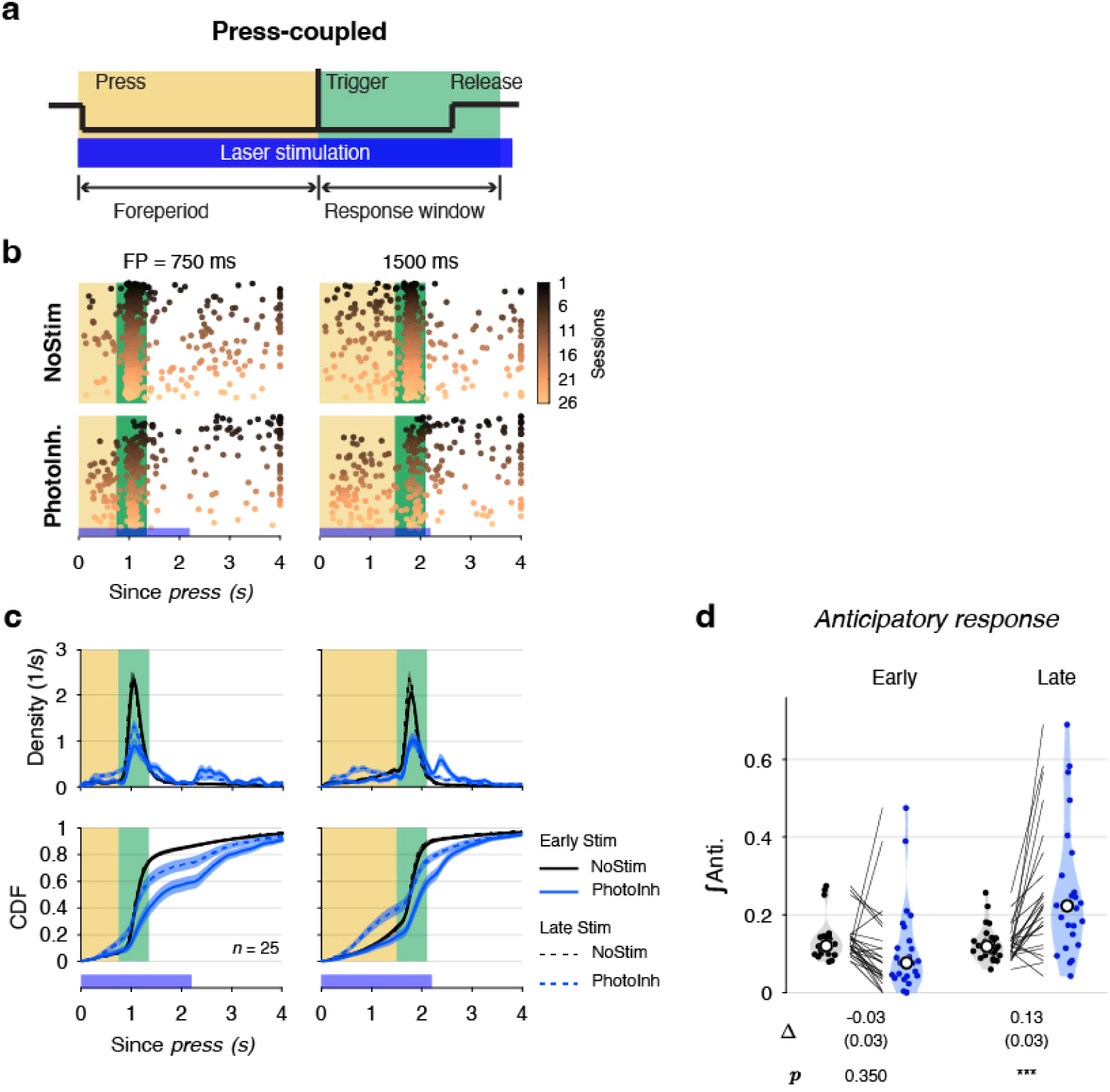
Press-coupled DLS photo-inhibition increases anticipatory responses. **a–c**, Similar to Fig. 6a–c, but for DLS photo-inhibition coupled to lever press and holding. **d**, Integral of anticipation function with (blue) or without (gray) photo-inhibition across *n*= 25 rats, grouped by photo-inhibition history (Early, Late). Mean differences (PhotoInh minus NoStim), SE, and Holm-adjusted *p*-values from pairwise EMM contrasts are displayed below. The anticipation function *G*(*t*) for premature responses was computed on 10-ms intervals from *t*= 0 to 1.5 s; the denominator of *G*(*t*) represents the number of trials with FP equals to or longer than the interval, and the numerator is the cumulative count of premature responses up to that interval:

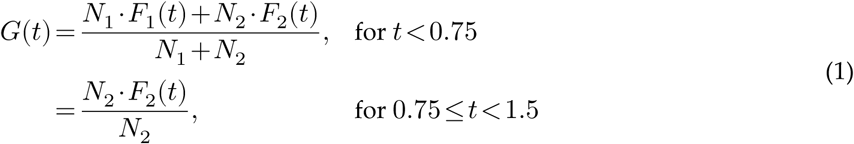

where *N*_*i*_ (*i*=1, 2) is the number of trials with FPs = 0.75 s and 1.5 s. *F*_*i*_(*t*) (*i*=1, 2) is the fraction of premature responses at or before time *t*. The integral (*y*) of *G*(*t*) from 0 to 1.5 s quantified premature response magnitude and was analyzed using an LME model to examine the effect of photo-inhibition on waiting (*y* ∼ StimType ×StimStage +(1+StimType ∣anm)):

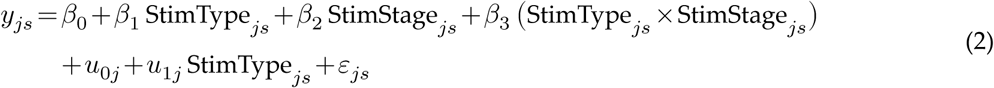

where *y*_*js*_ is the integral value of the anticipation function for StimStage *s* (Early or Late) in animal *j*.

**Fig. S10.**
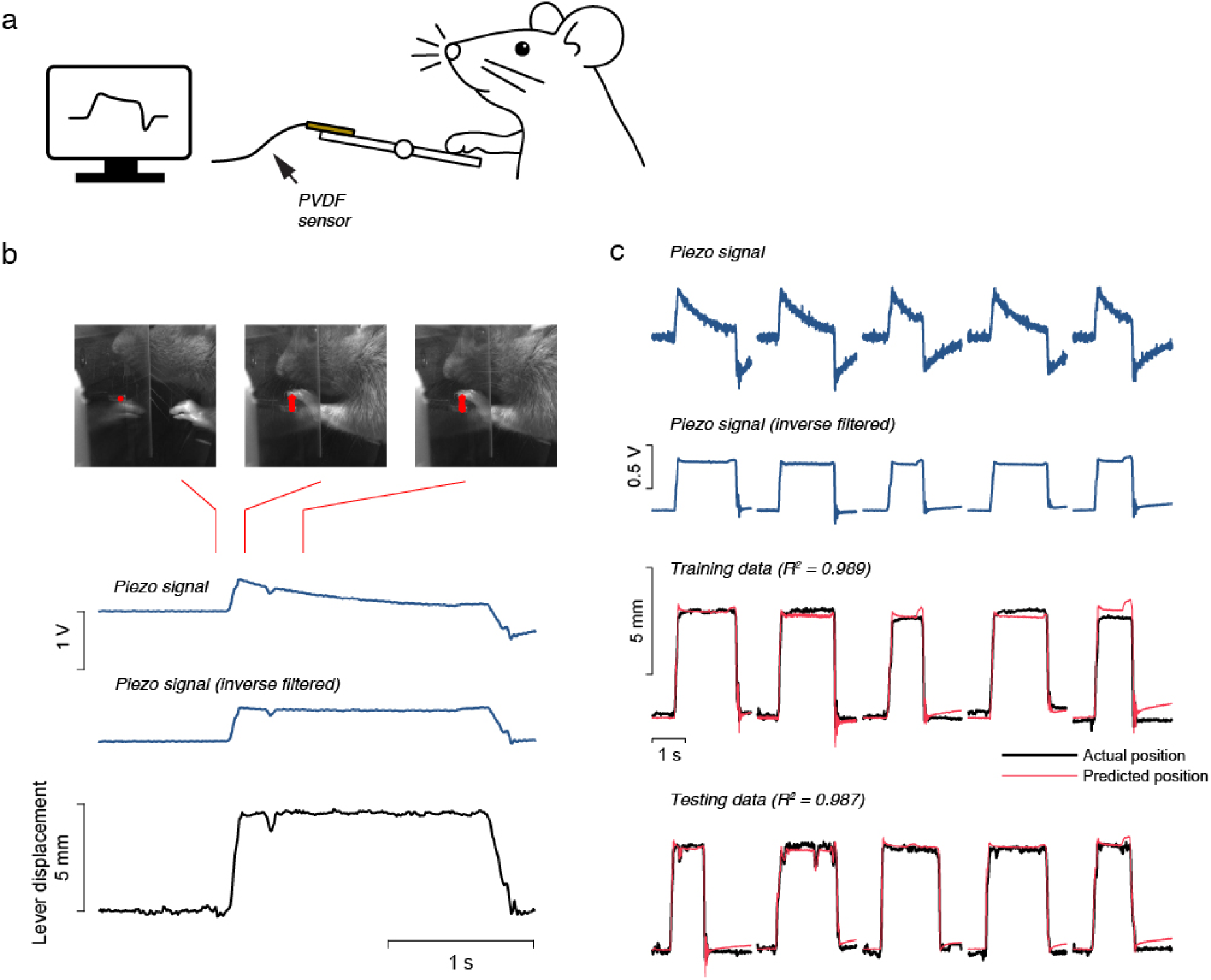
Piezo-sensor signal mapping to lever position. **a**, Diagram of a piezoelectric (PVDF) sensor attached to the opposite end of the lever to detect lever motion. **b**, Comparison of piezo-sensor signal and lever position during a lever press. The sensor signal, low-pass filtered during acquisition, was reconstructed via inverse filtering; lever position was tracked from video. **c**, Reconstructed piezo signal predicts lever position with a linear model, enabling rapid and accurate lever-position estimation.

---

**Movie S1.**
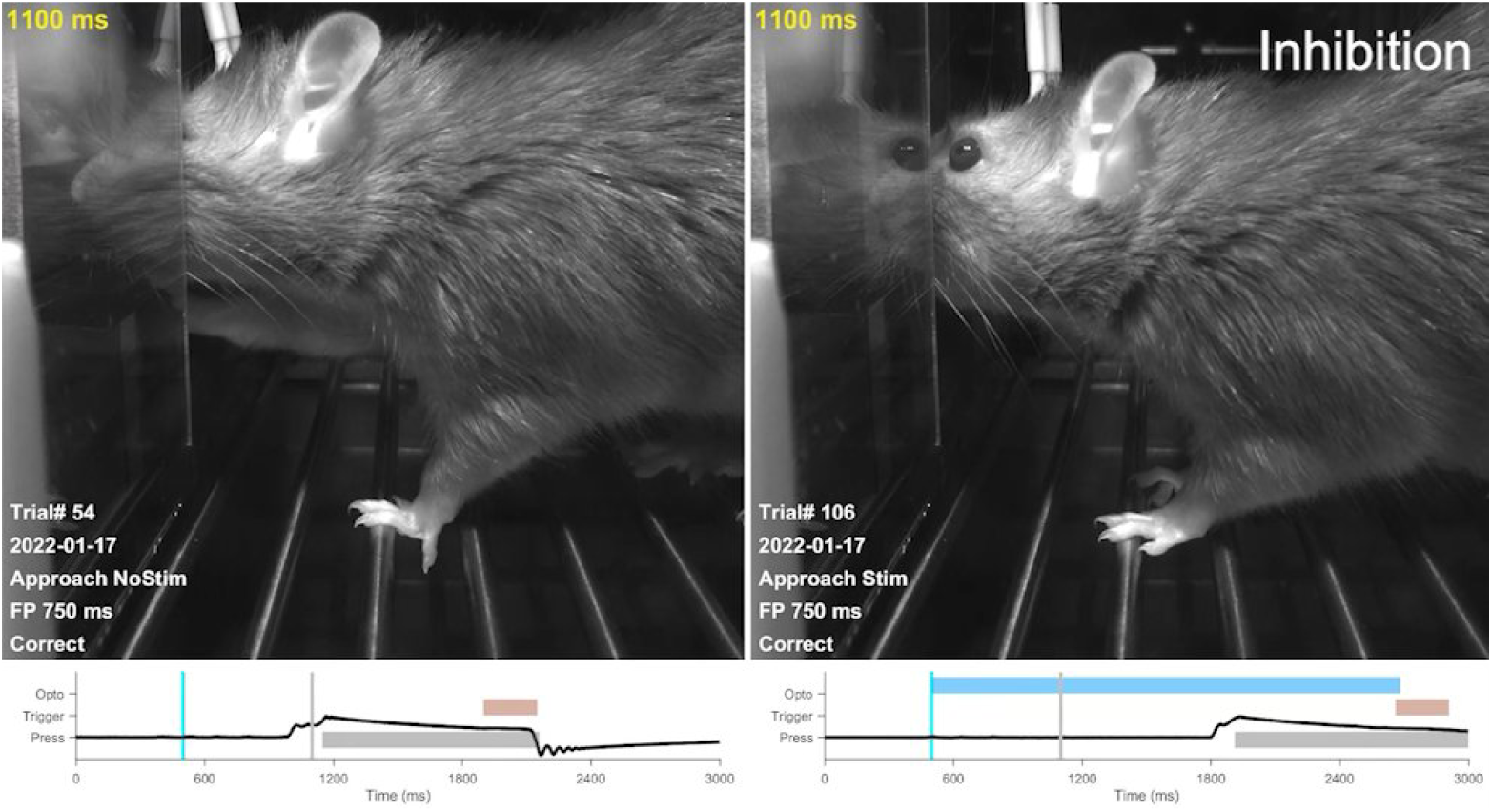
Comparison of forelimb reaching in a no-stimulation trial (left) and an approach-coupled DLS photo-inhibition trial (right). The movie plays at 0.1× real-time speed (tenfold slower). The bottom panel shows the piezo-sensor signal. Blue, brown, and gray shading indicate the duration of laser stimulation, trigger stimulus, and holding period. A cyan vertical line marks the approach detection in both panels. A gray vertical line marks the time of the current frame.

**Movie S2.**
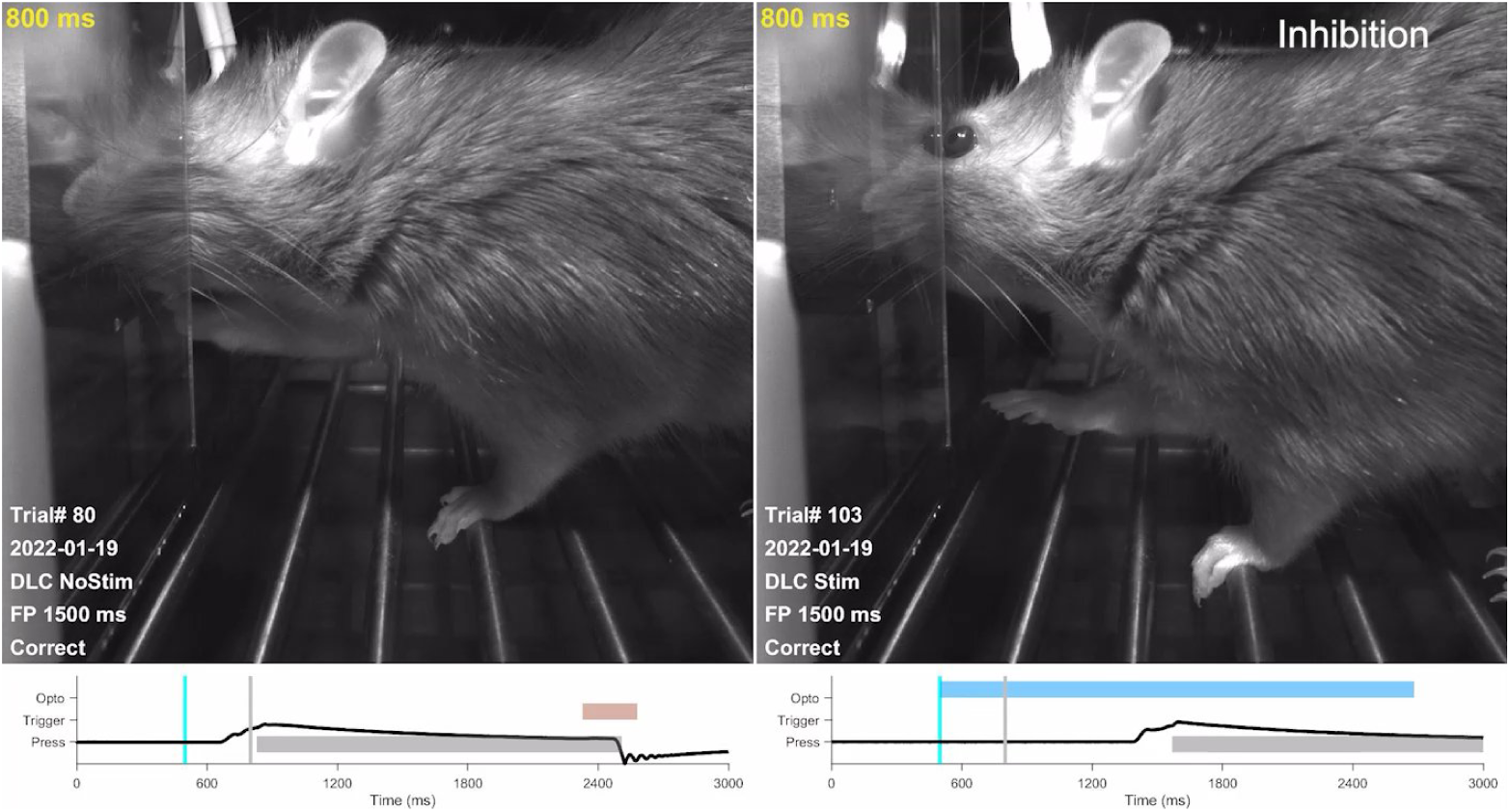
Comparison of forelimb reaching in a no-stimulation trial (left) and a lift-coupled DLS photo-inhibition trial (right). The movie plays at 0.1× real-time speed (tenfold slower). Panels match Movie S1; the bottom panel shows the piezo-sensor signal. Blue, brown, and gray shading denote the laser-stimulation period, the trigger stimulus, and the holding period, respectively. A cyan line marks paw-lift detection; a gray vertical line marks the time of the current frame.

**Movie S3.**
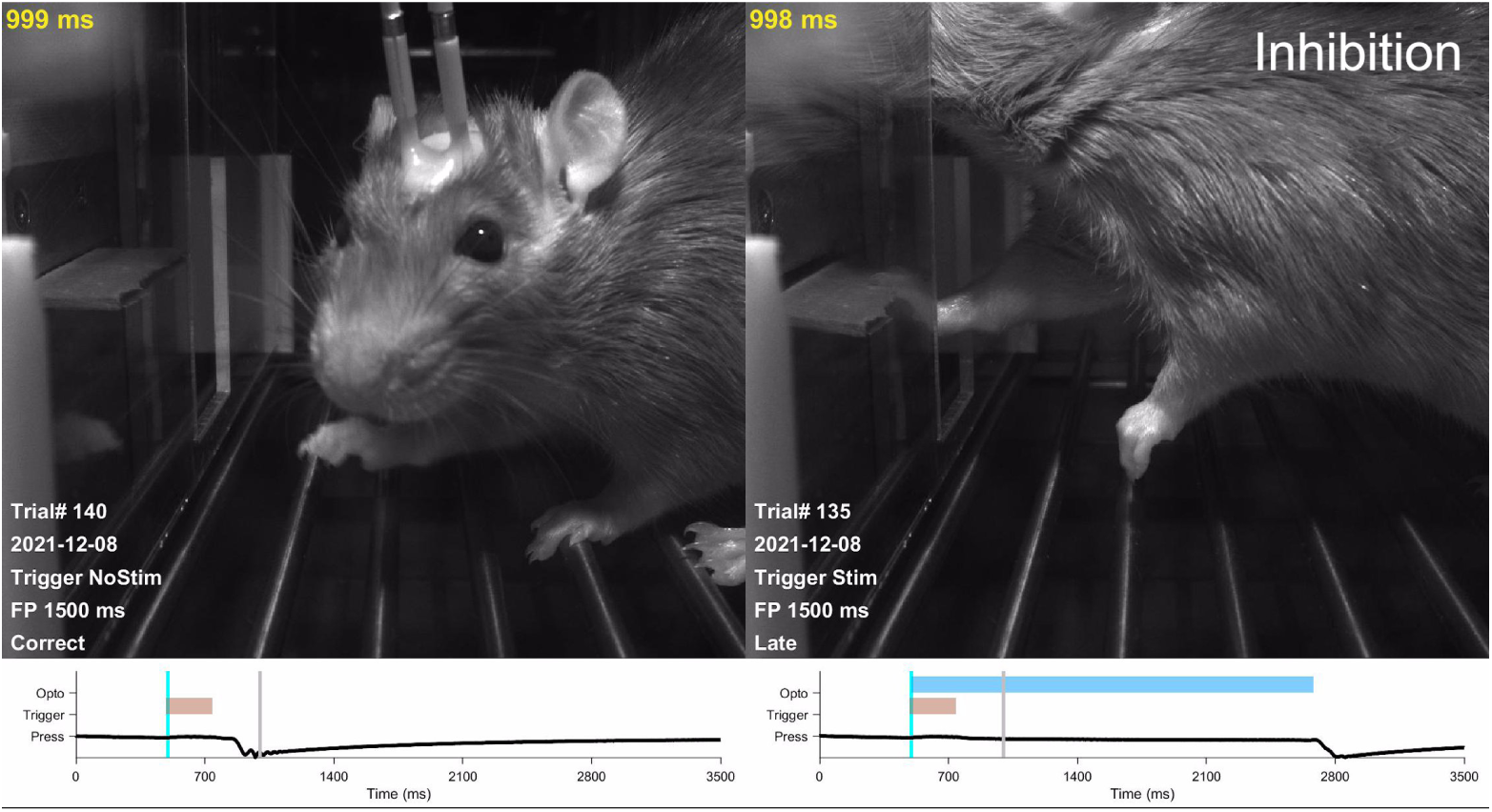
Comparison of lever release in a no-stimulation trial (left) and a trigger-coupled DLS photo-inhibition trial (right). The movie plays at 0.1× real-time speed (tenfold slower). Panels match Movie S1; the bottom panel shows the piezo-sensor signal. Blue, brown, and gray shading denote the laser-stimulation period, the trigger stimulus, and the holding period, respectively. A cyan line marks the trigger stimulus in both panels. A gray vertical line marks the time of the current frame.

**Movie S4.**
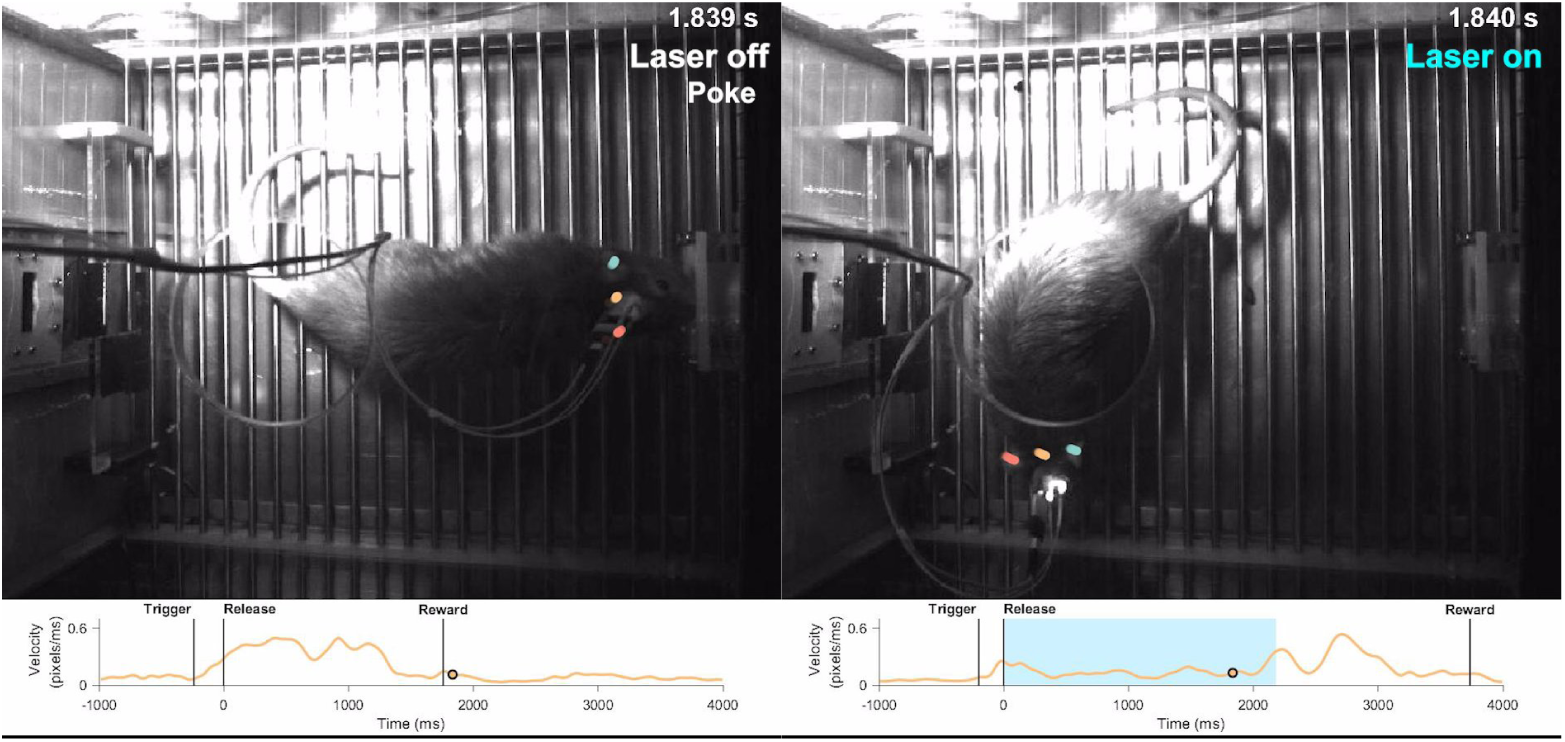
Comparison of reward retrieval in a no-stimulation trial (left) and a release-coupled DLS photo-inhibition trial (right). The movie plays at 0.2× real-time speed (fivefold slower). Tracking markers: red, right ear; green, left ear; yellow, midpoint between ears. The bottom panel shows the velocity of the midpoint. A circle tracks the time of the current frame.

**Movie S5.**
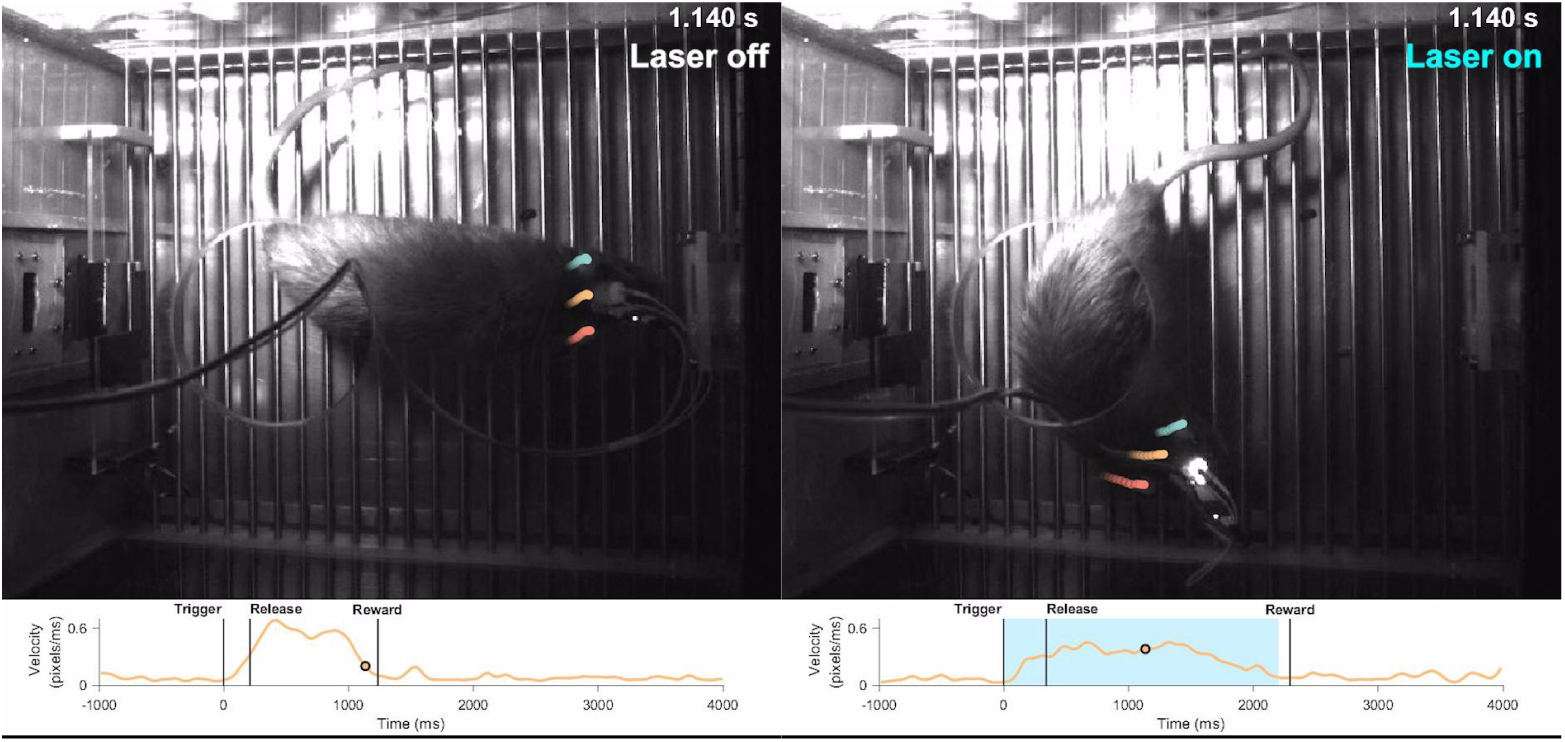
Comparison of reward retrieval in a no-stimulation trial (left) and a trigger-coupled DLS photo-inhibition trial (right). The movie plays at 0.2× real-time speed (fivefold slower). Tracking markers: red, right ear; green, left ear; yellow, midpoint between ears. The bottom panel shows the velocity of the midpoint. A circle tracks the time of the current frame.

**Movie S6.**
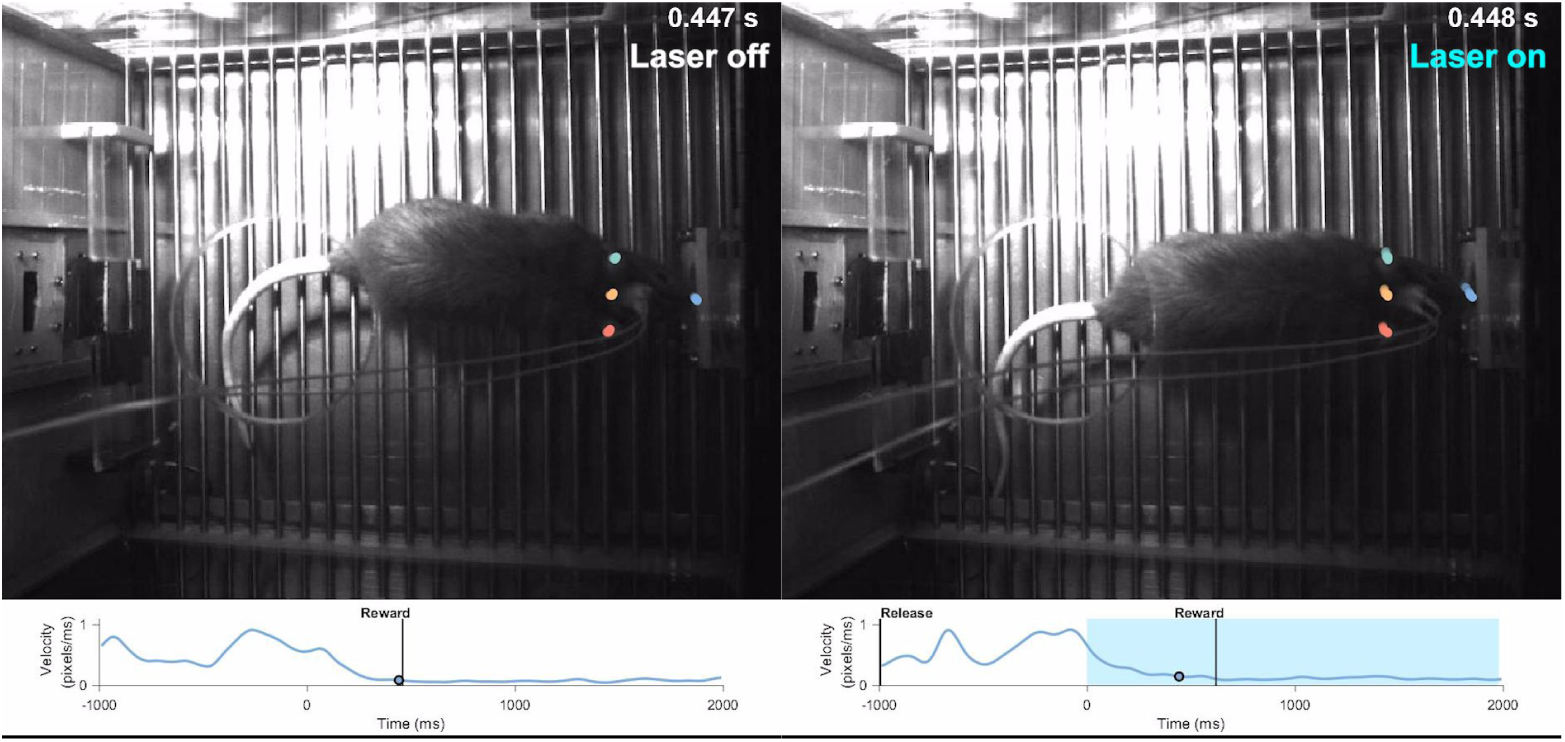
Comparison of reward retrieval in a no-stimulation trial (left) and a poke-coupled DLS photo-inhibition trial (right). The movie plays at 0.2× real-time speed (fivefold slower). Tracking markers: red, right ear; green, left ear; yellow, midpoint between ears; blue, nose. The bottom panel shows the velocity of the nose. A circle tracks the time of the current frame.

